# Learning *cis*-regulatory principles of ADAR-based RNA editing from CRISPR-mediated mutagenesis

**DOI:** 10.1101/840884

**Authors:** Xin Liu, Tao Sun, Anna Shcherbina, Qin Li, Kalli Kappel, Inga Jarmoskaite, Gokul Ramaswami, Rhiju Das, Anshul Kundaje, Jin Billy Li

## Abstract

Adenosine-to-inosine (A-to-I) RNA editing catalyzed by ADAR enzymes occurs in double-stranded RNAs (dsRNAs). How the RNA sequence and structure (i.e., the *cis*-regulation) determine the editing efficiency and specificity is poorly understood, despite a compelling need towards functional understanding of known editing events and transcriptome engineering of desired adenosines. We developed a CRISPR/Cas9-mediated saturation mutagenesis approach to generate comprehensive libraries of point mutations near an editing site and its editing complementary sequence (ECS) at the endogenous genomic locus. We used machine learning to integrate diverse RNA sequence features and computationally predicted structures to model editing levels measured by deep sequencing and identified *cis*-regulatory features of RNA editing. As proof-of-concept, we applied this integrative approach to three editing substrates. Our models explained over 70% of variation in editing levels. The models indicate that RNA sequence and structure features synergistically determine the editing levels. Our integrative approach can be broadly applied to any editing site towards the goal of deciphering the RNA editing code. It also provides guidance for designing and screening of antisense RNA sequences that form dsRNA duplex with the target transcript for ADAR-mediated transcriptome engineering.

## Introduction

RNA editing is an important mechanism that greatly diversifies the transcriptome and proteome in higher eukaryotes^1–5^. In animals, the predominant type of RNA editing is the hydrolytic deamination of adenosine (A) to form inosine (I), catalyzed by adenosine deaminase acting on RNA (ADAR)^6,7^. Abnormal A-to-I RNA editing is strongly linked to autoimmune diseases, neurological disorders and cancers^8,9^. Humans have two catalytically active ADAR proteins, ADAR1 and ADAR2, responsible for the editing of millions of RNA editing sites^10,11^. Adenosines in perfect or nearly perfect dsRNA duplexes, formed mainly by inverted repeats, are promiscuously edited^12^; while in non-repetitive sequences that often form imperfect dsRNA structures, ADARs can efficiently edit specific adenosines^13^. How RNA editing is regulated to determine its efficiency and specificity is poorly understood. It is proposed that both the primary sequence and secondary structure (*i.e.*, *cis*-acting regulatory elements) guide the preference and specificity of ADARs^7,14–19^. ADAR has a preferred sequence motif, in particular the 5’ and 3’ nearest neighboring positions (−1 and +1 nt) to the editing site^14–19^. Editing can be enhanced or suppressed by deviations from perfect base-pairing (*i.e.*, mismatches, bulges and loops), suggesting complex structural contributions to editing specificity^14–16^. The quantitative trait loci (QTL) mapping approach has been used to identify genetic variants associated with variability in RNA editing in *Drosophila* and humans, demonstrating that many editing QTLs (edQTL) can act through changes in the local and distal secondary structure for edited dsRNAs, consistent with importance of RNA structure^20,21^. Nevertheless, few general properties have emerged across different substrates, suggesting complex structural contributions to editing specificity^22^.

Previous studies are generally limited to small numbers of natural or engineered variants, thus lacking the systematic sequence and structure variation required for development of predictive models of editing, and a high-throughput, systematic mutagenesis approach is therefore called for. Predictive models derived through systematic mutagenesis would not only promote our understanding of the most prevalent type of RNA editing, but would also greatly advance the emerging field of ADAR-mediated transcriptome engineering, by informing the design of antisense RNA guides for therapeutic RNA editing. Prototypes of ADAR RNA guides were recently demonstrated to enable programmable RNA editing by forming dsRNA with target sites and recruiting endogenous ADAR proteins^23–29^. Improving the specificity and efficiency of programmable editing through rational design of guide/target duplexes will be critical as this new technology continues to be developed for clinical applications.

Here we combined CRISPR engineering, next-generation sequencing, and machine learning to decipher predictive *cis* regulatory RNA sequence and structural elements that affect the efficiency and specificity of ADAR1 mediated RNA editing. As proof-of-concept, we chose three representative RNA editing substrates and introduced hundreds of desired mutations at the endogenous loci in human cells using the CRISPR-mediated approach. We applied a supervised machine learning approach to build predictive models of substrate-specific RNA editing levels based on a variety of *cis* sequence and structural features. We identified general and idiosyncratic features that determine editing efficiency of individual substrates, highlighting the complexity of the *cis*-regulatory editing code. Our integrative approach, named predicting RNA editing using sequence and structure (PREUSS), lays the foundation for developing predictive models of RNA editing.

## Results

### CRISPR/Cas9-mediated mutagenesis to interrogate endogenous RNA editing

To interrogate the effects of *cis*-regulatory elements of RNA editing, we applied the CRISPR/Cas9 technology to introduce mutations at the endogenous loci of three known editing sites. The mutations were introduced both in the editing strand and the editing complementary sequence (ECS) that form dsRNA. Briefly, we designed guide RNAs (gRNAs) targeting regions of interest and oligonucleotide donors carrying mutations to directly knock-in mutations through the CRISPR/Cas9-mediated homology-directed repair (HDR) pathway^30^ (**Fig 1a**). To measure the RNA editing levels of the resulting variants, we performed targeted amplicon deep sequencing. Because the variant and the associated editing site are sequenced from the same read, there is no need to perform laborious clonal selection for homozygotes of the variants. Because each designed variant has a unique sequence, we successfully performed large scale multiplex mutagenesis and measured editing level without the aid of barcodes (see Methods for details).

**Figure 1:**
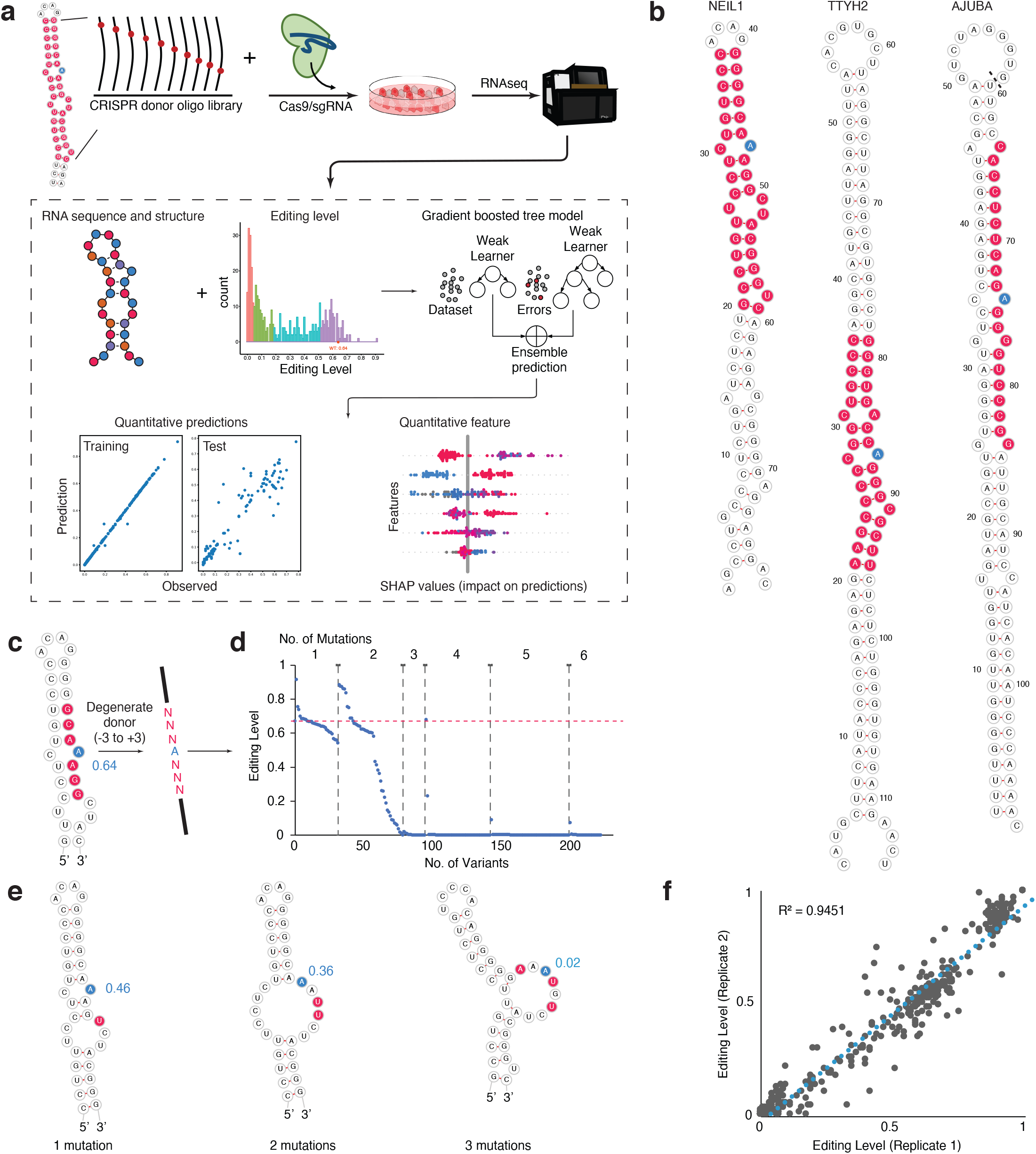
CRISPR/Cas9-mediated mutagenesis in endogenous RNA to dissect RNA editing by ADAR1 in cells. **a)** Overview of the experimental methods and computational pipeline. CRISPR/Cas9 mediated homology-directed repair is applied to mutagenesis of endogenous RNA in HEK293T cells. A supervised machine learning method (a gradient boosted tree, XGBoost) was applied to develop quantitative models that predict how cis elements such as RNA sequence and secondary structure determine RNA editing level. **b)** Sequence and secondary structure of the three RNAs, NEIL1, TTYH2 and AJUBA, for targeted mutagenesis. The residues subjected to mutations are highlighted in red and the specific editing site is in blue. For AJUBA, partial sequences from the genomic sequences are taken to focus on the region of interest. Therefore, the G59 and U60 shown in (**b**) is 524 nt apart in the genomic region. **c)** Degenerate donor oligos are designed for the −3 to +3 nt region around the specific editing site in the NEIL1 substrate. The mutagenized region is highlighted in red and the editing site in blue. **d)** The distribution of editing level by the number of mutations from the results of the degenerate NEIL1 library from (**c**). **e)** Examples of how the number of mutations affect the RNA secondary structure of NEIL1. **f)** Reproducible editing measurement of the two replicates of the targeted mutagenesis library of NEIL1.

We selected three natural ADAR1 substrates (NEIL1, TTYH2, AJUBA) (**Fig. 1b**) for the mutagenesis studies based on the observations that (1) the editing sites for all three substrates are highly edited (30−60%) in HEK293T cells, in which ADAR1 is expressed but ADAR2 is lowly expressed; (2) the editing sites are not edited when ADAR1 activity is abolished (data not shown); and (3) they represent three different types of dsRNA substrates. The NEIL1 editing site is in the coding region. The editing event leads to an amino acid change from Lysine (K) to Arginine (R), which has been shown to increase the enzymatic activity of the NEIL1 glycosylase^31^. The TTYH2 editing site is intronic and the AJUBA editing site is located in its 3’ UTR. The functional impact of these two editing sites is currently unknown.

In a pilot experiment of degenerate mutagenesis for NEIL1 and TTYH2, we randomized the region from −3 to +3 positions of the editing strand for NEIL1 (**Fig. 1c**) and a 10 nt region on the ECS of TTYH2 (**Supplementary Fig. 1a**). These degenerate mutations provide a rapid means to evaluate the CRISPR/Cas9 knockin efficiency and the effects of mutations. We observed that 3 or more mutations almost always lead to an abolishment of editing events (**Fig. 1d-1e, Supplementary Fig. 1b**). Therefore, to study variants that lead to a wide range of editing levels, we next performed targeted mutagenesis using a pool of 200 to 300 donors with designed mutations, focusing on single- and double-mutations with larger mutagenesis regions around the editing site in the editing strand and the ECS (**Fig. 1b, Fig. 2a, Supplementary Fig. 2a-2c**). For NEIL1 and TTYH2, we designed saturation single mutations both in the editing strand and the ECS (with the exception of –1 and +1 positions for NEIL where A to G mutations were eliminated as they would be indistinguishable from A-to-I editing). For AJUBA, we only designed mutations in the editing strand. We selectively designed double mutations (11% all possible double mutations) that can disrupt base-pairing at the mutation position. For NEIL1, because the ECS and editing site are close in sequence space, we were able to use donor oligos to introduce compensatory mutation variants which theoretically maintain base-pairing. Additionally, we designed indel variants to study the effects of selected secondary structure features of NEIL1, such as the bulge, internal loop, and stem length. Overall, between 10 and 20% of the sequenced RNAs carried mutations in the interrogated region, similar to previously reported knock-in efficiency^30^. We were able to reliably detect >90% of our designed variants after using stringent quality control filters. The knock-in results and editing measurements were highly reproducible (**Fig. 1f, Supplementary Fig. 1c-1k**). Interestingly, we discovered that using dsDNA as donor yielded similar results as using ssDNA oligonucleotides (**Supplementary Fig. 1j-1k**). Using dsDNA donors (e.g. PCR products) greatly simplifies the procedures and reduce the cost of the experiments.

**Figure 2:**
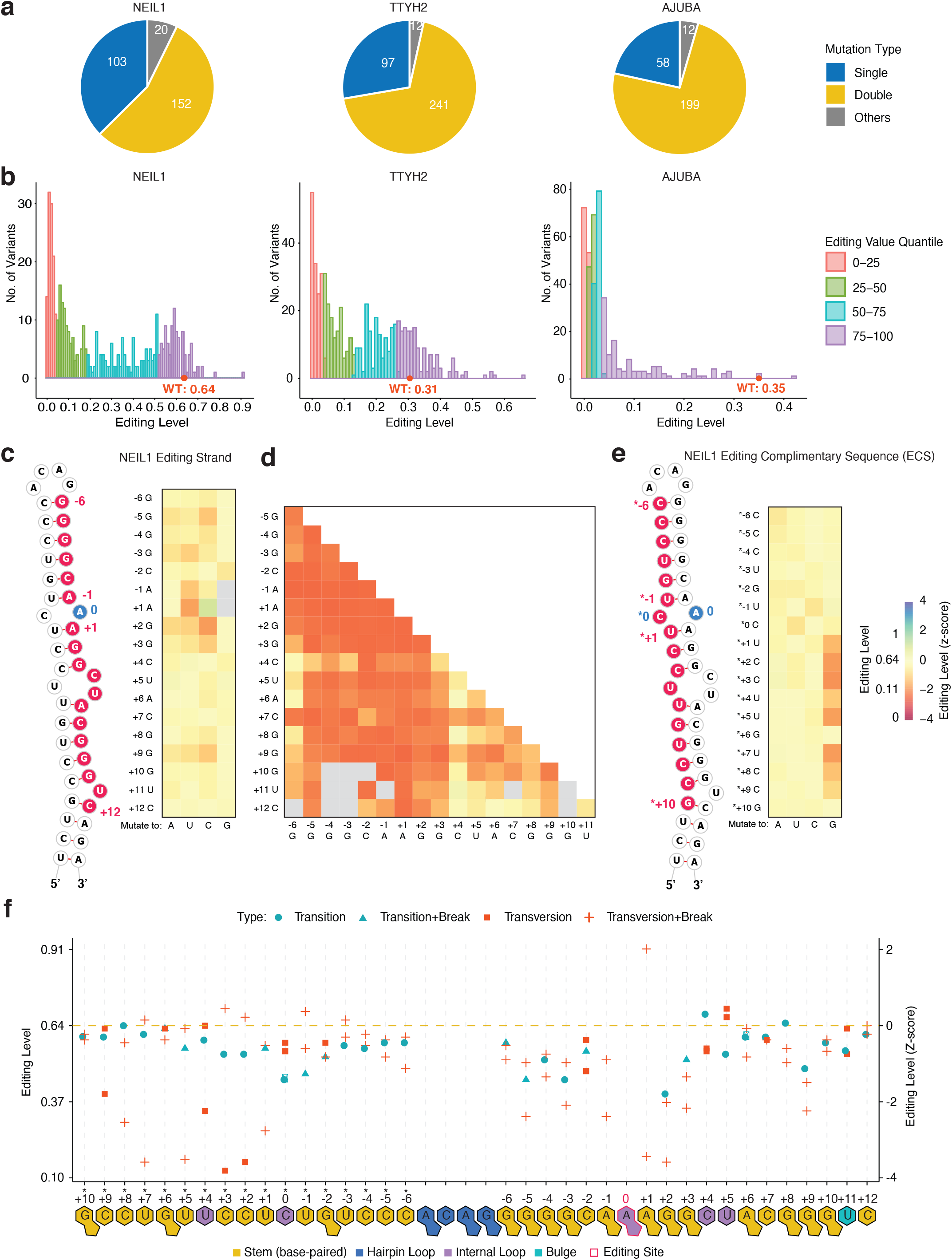
RNA editing results from the targeted mutagenesis experiments. **a)** Number of the types of mutations made in each targeted-mutagenesis library. **b)** Distributions of editing levels for each targeted-mutagenesis library, colored by editing level quantile in each RNA library. Heatmap of editing levels from **c)** single- and **d**) double-mutations in the editing strand of NEIL1. **e)** Heatmap of editing levels from single-mutation in the editing complementary sequence (ECS) of NEIL1. Editing level of WT NEIL1 is 0.64. The z-score is calculated as described in Methods and the WT editing level z-score is 0. c-e share the same heatmap scale shown in e). **f)** Position-specific effects of NEIL1 single mutations. The single mutations were grouped into four types: sequence change (transition and transversion) and the sequence change that also resulted in structural effects (break=disrupted the original WT base-pair at the position).

### Intertwined effects of primary sequence and secondary structure on editing levels

We compared the effects of single and double mutations in terms of the mutation type and mutation locations across all three RNA substrates (**Fig. 2, Supplementary Fig. 2**). We used computationally predicted secondary structure of each RNA variant to understand the associations between mutations and structure. We observed a wide range of effects from different variants of different substrates.

#### NEIL1

For NEIL1, most single mutations had minor effects on editing (−1 < *z*-score < 0), with the largest effects observed at positions +1 and +2 relative to the editing site (**Fig. 2c, Fig 3b**). Exceptions from this pattern were the large effects (*z*-score < −1) of G mutations downstream from the editing site in the ECS strand. This observation may suggest formation of alternative structures in these G-mutants (e.g. in **Fig. 3a**). Some RNA variants have the same predicted RNA structure but different editing level, suggesting a primary sequence effect (**Fig. 2b**). To decouple sequence and structural effects at each position, we also categorized the single mutations into four types (**Fig. 2f**). We observed that (1) there were large effects (*z*-score < −2) for the transversion mutations (change between purine and pyridine) that also disrupted base-pairing (transversion+break) at positions in close vicinity to the editing site (−5 to −1, +1 to +3, +9), and at the 3’ side of the editing site (*+1, *+5, *+7, *+8); and (2) at some positions (*+2, *+3) the primary sequence had larger effects than the disruption of base-pairing as demonstrated by larger effect of the transversion (*z*-score < −2) than the transversion+break mutations. Double mutations in the editing strand of NEIL1 had overall pronounced effects on editing (**Fig. 2d**). We observed different effects of these mutations according to their locations: (1) the strongest effects (*z*−score <− 2) were observed when at least one of the mutations was in close proximity to the editing site (positions –6 and +2); (2) the effect was generally smaller (*z*-score > −2) if one of the mutations was in a non-base-paired region (+4, +5, +11) (**Fig. 2d**).

**Figure 3:**
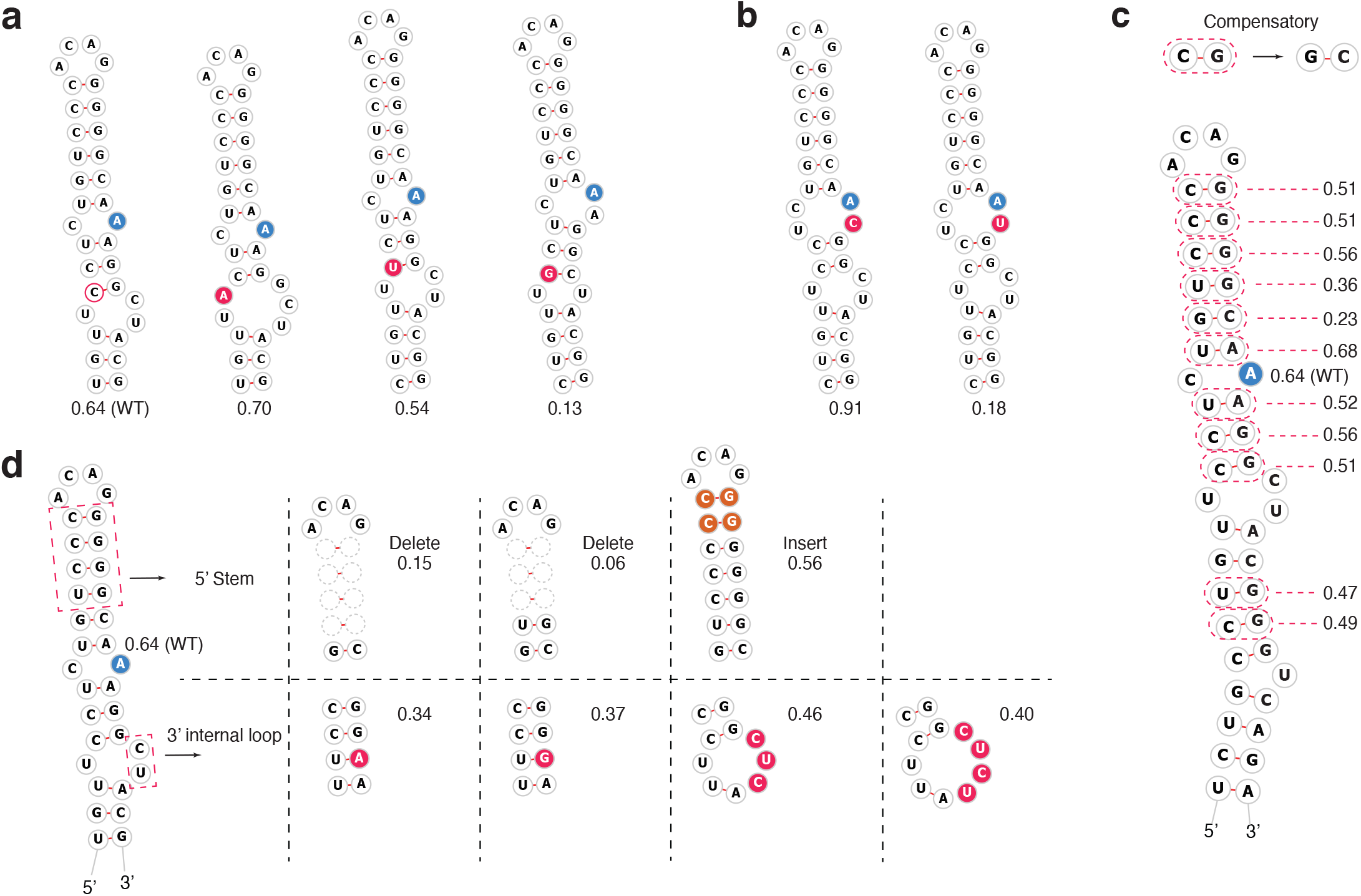
Examples of RNA secondary structure changes of NEIL1 variants. **a-b)** Single-mutation at different locations lead to different editing level. **c)** Compensatory mutations generally maintain high editing level. **d)** Alterations in the 5’ stem and 3’ non-stem structure elements affect editing level.

#### TTYH2

Contrary to NEIL1, a larger proportion of TTYH2 single mutations increased editing efficiency (*z*-score > 0) (**Supplementary Fig. 2a-2b**). TTYH2 has a lower WT editing level (30%) than NEIL1 (64%). Similar to NEIL1, the closer the single mutations were to the editing site (−2 to +1, *-3 to *+4), the larger their effects (*z*-score < −1) were on editing levels (**Supplementary Fig.3a**). Interestingly, several single mutations located both upstream (−6 to −3) and downstream (+6 to +8, *+6) of the editing site increased editing levels (*z*-score > 0). For TTYH2 double mutations, the effects were most negative when at least one mutation was located in –2 to +6 in the editing strand and *−3 to *+5 in the ECS.

#### AJUBA

Editing level of WT AJUBA is 35%. In contrast to NEIL1 and TTYH2, many single mutations of AJUBA were sufficient to disrupt editing. Also, all double mutations abolished editing regardless of positions (**Supplementary Fig. 2c**). Most of the transversion single mutations disrupted base-pairing and the transversion+break mutations had larger effects (z-score < −1) when located closer to the editing site (**Supplementary Fig. 3b**). Contrary to NEIL1 and TTYH2, the AJUBA single mutations with largest effects (z-score < −2) were transitions (change within purine or pyrimidine) and transition+breaks, suggesting that primary sequence might have a larger influence on editing of the AJUBA RNA. This difference between editing targets may be explained by much lower predicted thermodynamic stability of the AJUBA secondary structure (free energy of −71 kcal/mol for AJUBA vs. −87 kcal/mol for TTYH2 of similar length), which would make the relative effects of a mismatch in AJUBA larger than those in RNAs with more stable structures.

Taken together, these observations indicate that RNA primary sequence and structure are intertwined in terms of mediating effects of mutations on editing efficiency for different substrates. These effects also vary among the three different RNA substrates, suggesting substrate-specific *cis*-regulation rules.

### Structural features of the editing site, 5’ stem and 3’ non-stem elements, and RNA stability affect editing levels

Next, we systematically explored the effects of changes to secondary structure on RNA editing levels. We found that compensatory mutations in NEIL1 that did not affect secondary structure resulted in only minor reduction of editing levels (**Fig. 3c**), implying that mutations may need to impact secondary structure to have major effects on RNA editing efficiency. To further investigate how a specific structural feature affects editing efficiency, we designed several indels to alter secondary structures of NEIL1 (**Fig. 3d**). The 5’ stem length is critical for NEIL1, as both shortening it and breaking the base-pairing abolished editing, and increasing the stem length by 2-bp did not increase editing efficiency either (**Fig. 3d, Supplementary Fig. 3c**). The 3’ base-pairing is also critical because breaking it led to nearly complete disruption of editing (**Supplementary Fig. 3c**). When we replaced the downstream 3’ internal loop with either a canonical base-pair or wobble base-pair, the editing efficiency decreased by ~50% (**Fig. 3d, Supplementary Fig. 3c**). Enlarging the loop with additional nucleotides resulted in mild (−1< z-score <0) reductions in editing levels (**Fig. 3d**). These data suggest that a duplexed 5’ stem is important for ADAR1 editing and a non-stem feature downstream at the 3’ is also required for efficient editing. Additionally, editing site structures containing an A:C mismatch (also named as a 1:1 internal loop) exhibited higher editing levels on average than when the editing site resided in a larger loop (*P* < 0.0001, **Supplementary Fig. 3d**), consistent with previous structural and biochemical studies^29^. However, several editing site structures harboring non-A:C mismatches also showed strong editing levels for NEIL1 and TTYH2 (Fig. 3b **and Supplementary Fig. 3d)**, indicating that additional factors affect editing efficiency.

We reasoned that features affecting RNA thermodynamics could also affect editing efficiency^18^. We observed significant differences in the Free Energy (**Fig. 4a**) and All Stem Length (**Fig. 4b**) between highly (highest 25 percentile of editing level in each RNA library) and lowly (lowest 25 percentile) edited NEIL1 (*P* < 0.0001) and AJUBA (*P* < 0.001) variants, suggesting RNA stability is important in determining editing activity for these two RNA substrates. No statistically significant difference was observed for TTYH2 variants. We also hypothesized that RNA variants that are structurally more similar to the WT would result in editing levels similar to WT. We quantified structural similarity using two measures: “Similarity Score” that indicates the degree to which the structure of a variant is similar to the structure of the WT and “Probability of Active Conformation” that indicates the probability of forming wild type secondary structure in the structure ensemble in each RNA variant. We found significant differences (*P* < 0.0001) of similarity score between highly and lowly editing variants only for NEIL1 but not for the other two substrates (**Fig. 4c**). A higher probability of active conformation was observed in the highly edited variants compared to lowly edited variants for all three substrates (*P* < 0.0001, **Fig. 4d**). Taken these two results, the WT structure is not necessarily the only active but one of the more preferred conformations. However, when we considered all variants across the entire editing spectrum, instead of highly vs. lowly edited variants, no significant correlations were observed between these features and RNA editing levels (**Supplementary Fig. 4**). These results show that individual sequence, structure and stability features of variants only have limited predictive association with quantitative editing levels. Therefore, we decided to integrate multiple RNA sequence and structure features to model quantitative editing levels via machine learning.

**Figure 4:**
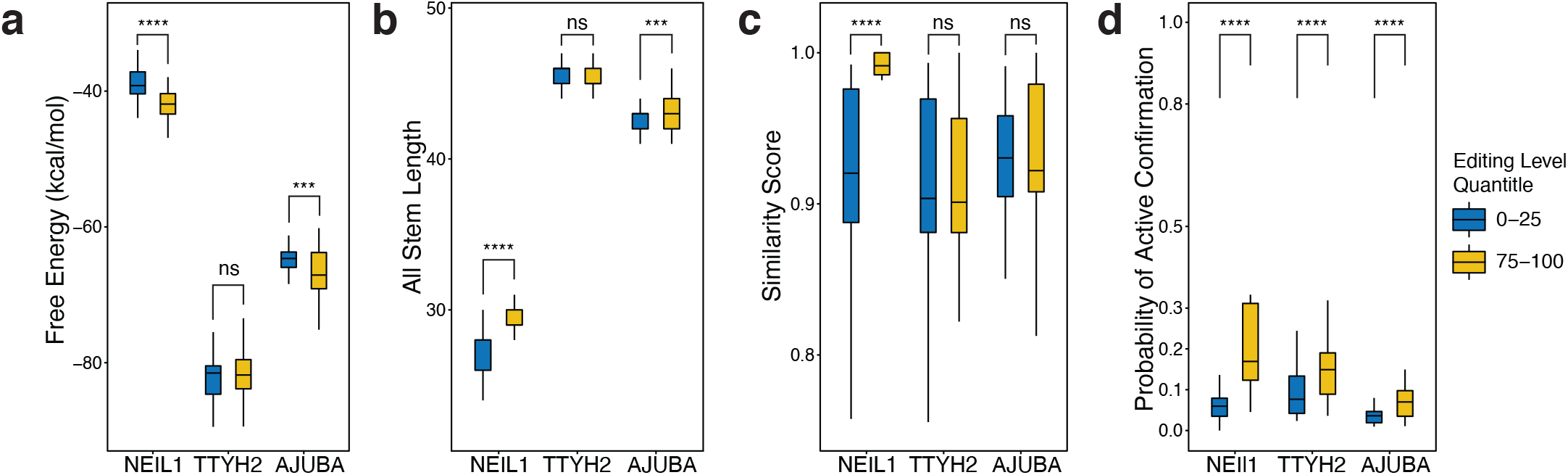
*Cis* regulatory features explain differences of editing levels among RNA variants. Comparing the difference of **a)** Free Energy, **b)** All Stem Length, **c)** Similarity Score, and **d)** Probability of Active Conformation in the highly edited (75 to 100 percentile in editing level in the library) with the lowly edited (0 to 25 percentile) variants in each RNA library. ****, *P* < 0.0001 in t-test.

### Machine learning models accurately predict substrate-specific RNA editing levels from sequence and structure features

Given the observation that no single property of the RNA substrates correlated strongly and consistently with editing efficiency, we turned to machine learning models to capture the complex relationship between editing levels and multiple RNA sequence and structure features. A set of 122 features were derived to annotate the RNA variants (see **Methods** and **Supplementary Tables 2-4** for feature annotations for all variants of NEIL1, TTYH2, and AJUBA respectively). The sequence features summarized various properties of the primary RNA sequence of each variant at and around the vicinity of the editing site. We used the bpRNA^32^ tool to assign all residues in each variant to diverse structural elements such as hairpin loops, bulges, internal loops, stems, multi-loops and closing pairs (**Fig. 5a**). Structure features were derived at the editing site and adjacent regions up to 3 bpRNA structural elements upstream and downstream from the editing site (**Fig. 5b**), as these regions within the RNA substrate fully encompass the interaction site with the ADAR deaminase domain (**Fig. 5c**)^33,34^. The 122 features were further grouped into nine major categories for purposes of feature interpretation (**Supplementary Fig. 1**). Gradient boosted trees (GBTs) were trained via the XGBoost algorithm^35^. We trained and tuned GBTs on distinct subsets of RNA variants to map their feature annotations to corresponding real-valued editing levels or binarized labels obtained by thresholding editing levels into two classes (edited vs. not-edited).

**Figure 5:**
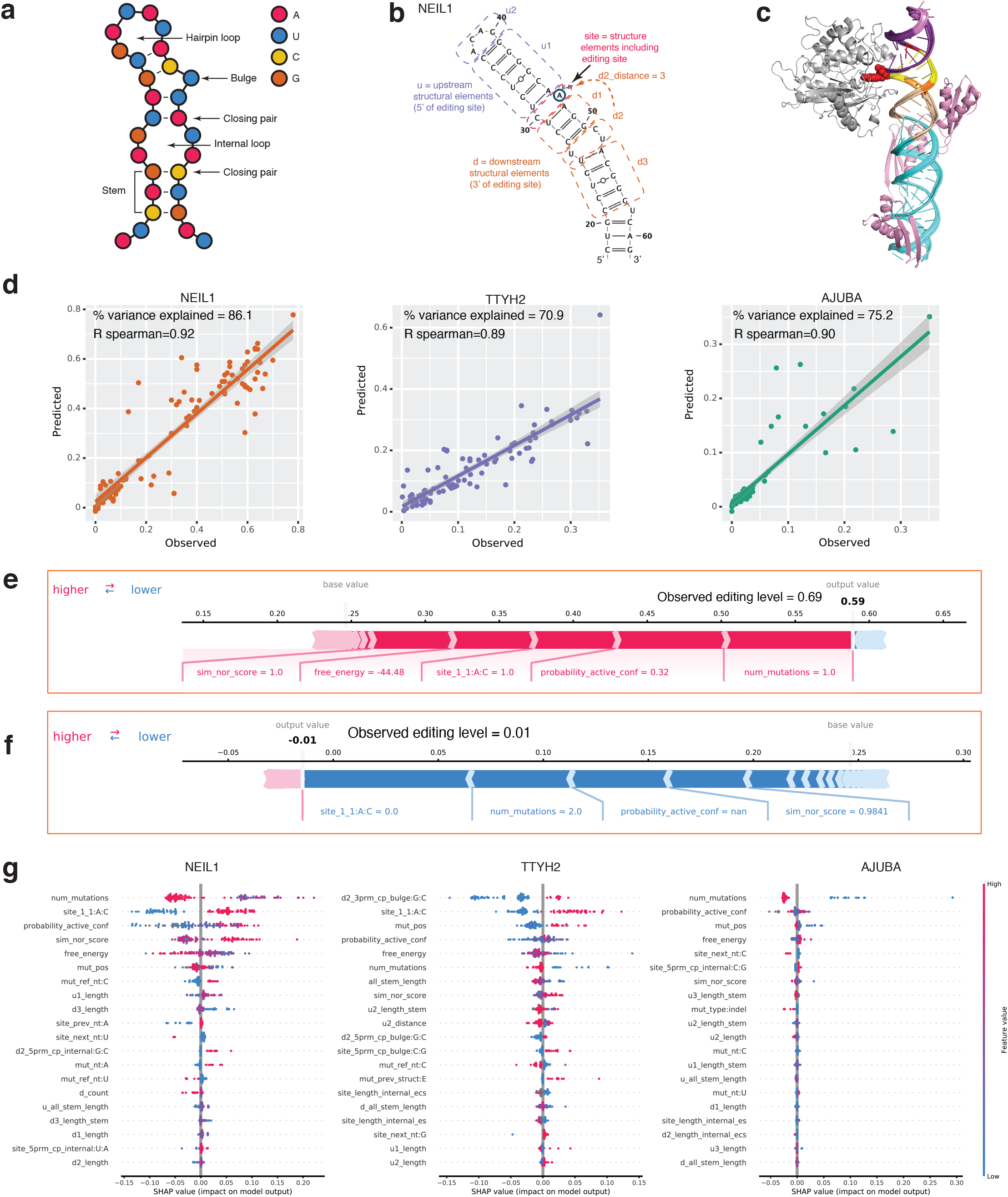
Quantitative model predicts editing level by combining complex RNA sequence and structure features. **a)**Structural features annotated by bpRNA and included in featurization of RNA variants. **b)** High-level feature groups for input to XGBoost analysis. u1= structural element immediately upstream of editing site; u2= structural element upstream of u1; site=structural element within which the editing site is found; d1= structural element downstream of site; d2= structural element downstream of d1; d3= structural element downstream of d2. **c)** Illustration of a putative model for binding of the NEIL1 RNA (cyan) to the ADAR1. The ADAR1 deaminase domain (silver) and modeled from ADAR2 by Phyre2. The dsRNA binding domains (pink) are modeled in one possible conformation as described in the Methods. The editing site 1:1 internal loop on NEIL1 is shown in red and the editing A shown as space filled. The upstream (purple and light purple) and downstream (yellow, orange and light orange) immediately adjacent to the editing site are colored according to shown in (b). **d)** XGBoost editing level predictions for variants of NEIL1 (orange), TTYH2 (purple), and AJUBA (green) within the test split (15% random split of positions). *R*^2^ is a measure of the % variance explained. Spearman R indicates correlation between observed and predicting editing values. **e)** SHAP annotation of feature contributions for the NEIL1 test split variant with the highest observed editing level. Features with positive SHAP scores (drive the prediction over the dataset base value) are indicated in pink; features with negative SHAP values (drive the prediction below the dataset base value) are indicated in blue. Base value refers to the mean predicted editing level across the test split. Output value refers to the XGBoost prediction on this example. The four features with the highest absolute value SHAP scores are shown. **f)** SHAP annotation of feature contributions for the NEIL1l1 test split variant with the lowest observed editing level. **g** SHAP values for the 20 most important features driving XGoost editing level predictions on the test split for NEIL1, TTYH2, and AJUBA. Each dot indicates a variant in the test split. Features (y-axis) are ranked from top (most significant) to bottom (least significant) by predictive importance.

First, we evaluated the prediction performance of our model for each substrate. We trained and tuned models on a subset of variants and then tested model performance on a held-out test set of variants of the same substrate. For NEIL1, the models accounted for 86% of the variance (*R*^2^) in ADAR editing levels for variants in the held-out test set, with a Spearman correlation (*R*_*s*_) of 0.91 between observed and predicted editing levels. Binary editing status was also predicted accurately (Area under precision-recall curve, *auPR* = 0.99). Similarly, high test set predictive performance was obtained for TTYH2 variants (*R*^2^ = 0.71, *R*_*s*_ = 0.89, *auPR* = 0.98) and AJUBA variants (*R*^2^ = 0.75, *R*_*s*_ = 0.90, *auPR* = 0.96). Augmenting the training set for each substrate with variants from the other substrates did not result in any significant improvements in model performance (**Supplementary Table 5, Supplementary Fig. 6**). These results indicate that it is possible to predict RNA editing levels of new mutations in a substrate with high accuracy from sequence and structure features using integrative machine learning models trained on a subset of mutations from the same substrate (**Fig. 5c**).

Next, we tested whether models trained on variants of one or more substrate could predict editing effects of mutations in a different substrate. We observed a significant drop in model performance for cross-substrate prediction of RNA editing (**Supplementary Fig. 6, Supplementary Table 8**). For example, a model trained on NEIL1 variants yielded lower performance on AJUBA variants (*R*^2^ < 0.05, *R*_*s*_ = 0.56, *auPR* = 0.82) and TTYH2 variants (*R*^2^ < 0.05, *R*_*s*_ = 0.51, *auPR* = 0.65) as compared to a model trained and tested on NEIL1 variants (*R*^2^ = 0.86, *R*_*s*_ = 0.91, *auPR* = 0.99). Similarly, a model trained on all NEIL1 and TTYH2 variants also yielded lower predictive performance when tested on AJUBA variants (*R*^2^ = 0.22, *R*_*s*_ = 0.53, *auPR* = 0.86), albeit higher than the model trained on either substrate independently. The same held true for all models trained on two of the substrates and evaluated on the third – lower performance compared to within-substrate training and evaluation. An attempt to normalize numerical feature values for variants of all three substrates relative to the wild type (WT) did not significantly improve the model’s performance on the test sets **(Supplementary Table 8, Methods**).

The inability of our current models to accurately generalize predictions to new substrates is not entirely surprising considering the diversity of the substrates and the small number (three) of distinct substrates available for model training. It is likely that the challenge of cross-substrate training may be solved by training on a larger number of variants from a database of diverse substrates, and future efforts will focus on this task. However, given the success of our substrate-specific models in predicting editing effects for unseen mutations within each substrate, we decided to interpret these models to investigate the features that may be predictive of RNA editing levels.

### Model interpretation provides insights into common and substrate-specific features associated with RNA editing efficiency

For each of the three substrate-specific models, we used the TreeExplainer SHAP (SHapley Additive exPlanations) algorithm to quantify the contributions (or importance) of all features to the RNA editing predictions of each variant in the test sets^35^. The SHAP importance score of a feature with a specific value for a variant of a substrate estimates how much the feature contributes to pushing the model’s output from a baseline editing level to the predicted RNA editing level for the variant. The baseline editing level is defined as the average editing level across all variants in the test set. Examples of how SHAP scores illuminate feature importance are illustrated in **Fig. 5d and 5e. Figure 5d** illustrates the SHAP scores for the 5 most important features for the NEIL1 test variant (RNA ID 92, 31UtoG) with the highest observed editing level (0.69). The NEIL1 model predicts an editing level of 0.59 for this variant. For the NEIL1 test set, the baseline (mean) predicted editing value is 0.25. We display the contribution of all feature values for this variant in pushing the prediction from the baseline of 0.25 to the predicted output value of 0.59. The feature “num_mutations=1”, which encodes the number of mutations in the RNA equal to 1, is estimated to have the highest importance and increases the prediction by 0.09 from the baseline. The contribution of the editing site taking on an A:C mismatch pushes the prediction up by approximately 0.05, and so on. Conversely, **Figure 5e** illustrates how feature values unfavorable to ADAR editing result in a predicted editing level of −0.01 for another variant (ID 142, 41GtoC,45CtoG) of the NEIL1 substrate relative to the baseline level of 0.25. This variant has two mutations in the substrate (num_mutations=2). This feature has a SHAP score of −0.05, indicating that a higher number of mutations in the RNA is unfavorable to editing. There is no A:C mismatch at the editing site, and this feature value has a SHAP score of −0.06. These and other highlighted feature values serve to drive the prediction down by 0.26 from the baseline value.

To understand the directionality of predictive association of the features with RNA editing levels, we plotted the SHAP scores of the top 20 features for all test-set variants of the three substrates (**Fig. 5f**). We also summarized the relative importance of features for each of the three substrates by computing the percent contribution from each feature to the mean of absolute SHAP values across all examples in the test sets of each substrate (**Fig. 6a**).

**Figure 6:**
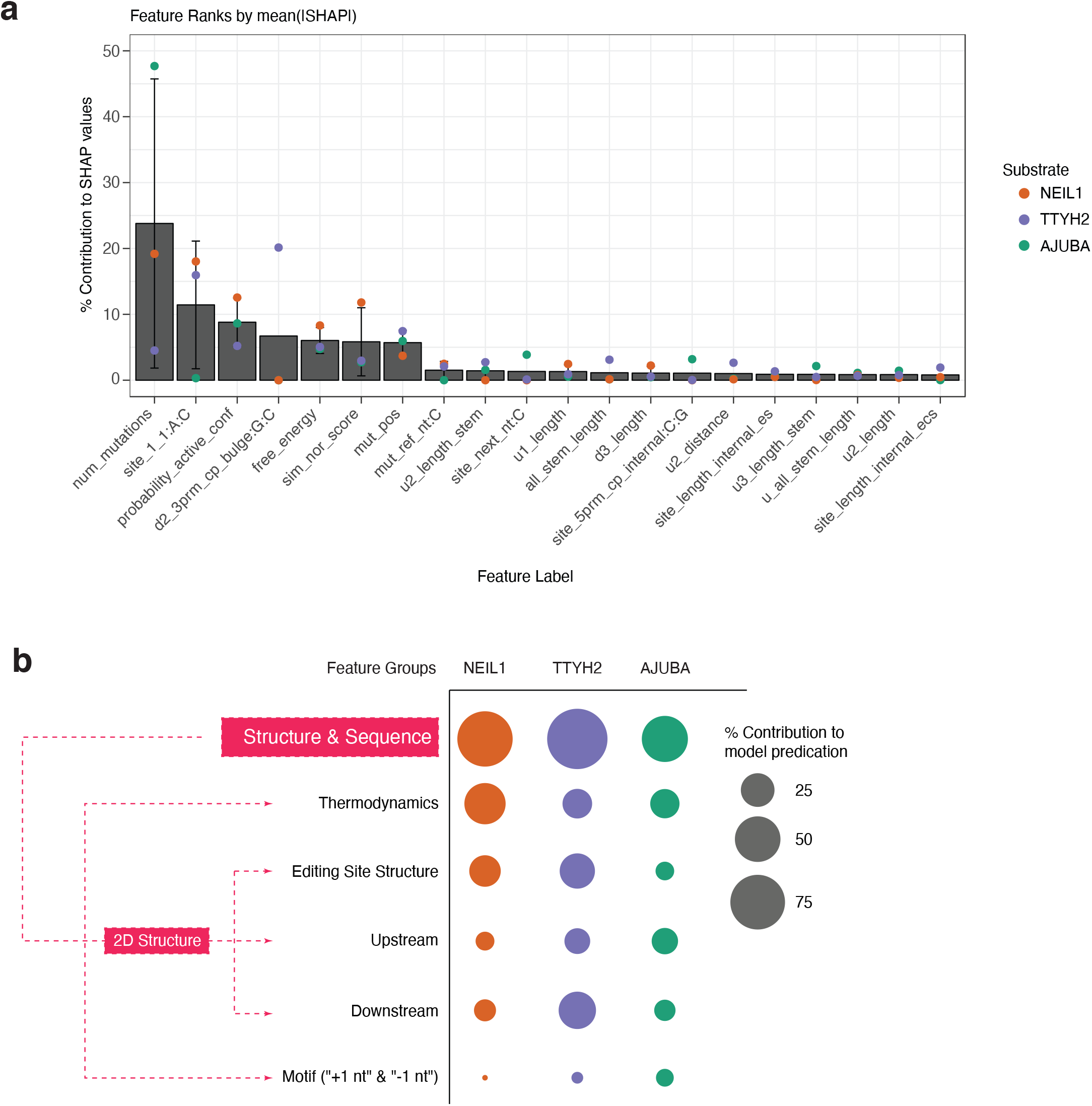
*Cis* regulatory features synergistically contribute to model prediction. **a)** Percent contributions of individual features to model prediction ranked by averaging normalized SHAP values. Higher ranking with smaller standard errors indicates that these features are commonly among the highest contributor to model prediction in all three RNAs. **b)** Contributions of different feature groups to the prediction of editing levels for each RNA library. The group of individual features included in each feature group are listed in Supplementary Table 1.

Seven features were illuminated as most important for driving model predictions across substrates. Number of mutations (“num_mutations”) was the strongest contributor for AJUBA (47.69%) and NEIL1 (19.18%) and in the top 6 most important features for TTYH2 (4.51%) (**Fig. 6a**). Increasing number of mutations had a negative influence on editing levels (**Fig. 5f**). This effect supports the proposal that RNA structure plays a big role in editing activity because in our library design the more mutations (single vs double) the larger changes occur in the structure (**Supplementary Fig. 2e**). An A:C mismatch at the editing site (“site_1_1:A:C”) had a high relative contribution for NEIL1 (18.03%) and TTYH2 (15.95%) but contributed less to AJUBA editing levels (0.31%). The presence of A:C mismatch was strongly positively correlated with editing levels, consistent with previous proposals that it facilitates the flipping-out of the adenosine for ADAR editing^33,34^ (**Fig. 5f**). The probability of the active conformation (“probability_active_conf”) accounted for a mean of 8.8% relative contribution across substrates. A G:C closing pair at the 3’ end of a bulge structure downstream of the editing site (“d2_3prm_cp_bulge:G:C”) was highly specific to the TTYH2 substrate, with the strongest relative contribution of 20.14% for that substrate. However, this feature was not present in the NEIL1 and AJUBA substrates, as the d2 structure in these was not a bulge. More negative free energy (“free_energy”) had a positive influence on editing levels **(Fig. 5f)**. The structure-similarity score of variants compared to WT feature (“sim_nor_score”) was positively correlated with editing levels. These four features corroborate with previous results that the overall structural stability of RNA substrate is positively correlated with the editing activity^18^. The seventh ranking feature was the position of the mutation along the RNA molecule (“mut_pos”). “Mut_pos” values are numbered beginning at the 5’ end of the RNA molecule, so higher values indicate positions further from 5’ and closer to 3’. This result indicates that the nucleotides adjacent to the editing site in the structure is the hot spot dictating activity. Though the “mut_pos” feature had a strong impact on editing level, the directionality varied across substrates, reflecting the interplay of the mutation position with other structural features.

In addition to top individual features, a sparse set of features collectively contribute to the accurate predictions made by the models. For the NEIL1 substrate, 90% of the explained variance could be attributed to the 26 top features, compared with the 32 top features for TTYH2 and 23 top features for AJUBA (**Supplementary Table 9**). To illustrate the contributions of different types of features and to draw biological insights, we looked at feature groups and subgroups. We categorized the group of all structure and sequence features excluding the ones related to mutation-related features to 4 subgroups (**Fig. 6b**, full list of feature groups and subgroups in **Supplementary Table 1**). Overall, the thermodynamics and the editing site structure have the largest contributions, consistent with prior proposals that the overall RNA stability and the structure of the editing site dictate the editing efficiency^14,18^. Notably, upstream and downstream structure features are also important, such as the downstream features in TTYH2. The −1 and +1 nt sequence motif (termed site_prev_nt” and “site_next_nt” in **Fig. 5g** and **Supplement Table 1**) also contributes to the prediction albeit to a lesser extent. This is likely due to the fact that the motif preference was identified from previous analysis on well-controlled RNA secondary structure^14^, while our approach encompasses comprehensive sequence and structure features.

This systematic interpretation of our models not only reveal several biologically relevant features that are globally predictive across the three substrates, but also some that are highly predictive for specific substrates. These results showcase the promise of predictive *cis*-regulatory models of RNA editing but also highlight the need for much larger datasets spanning diverse substrates to learn more generalizable models of RNA editing.

## Discussion

The ultimate goal in understanding the *cis*-regulation of RNA editing is to develop a model that accurately predicts the ADAR editing efficiency *in vivo*, namely an “editing code”. Unlike protein-DNA or protein-ssRNA interactions, where the primary *cis*-sequence largely dictates the interaction, ADAR substrates are required to bear double-stranded secondary structure. The difficulty of associating RNA sequence and secondary structure features to editing activity is a major challenge in studying the *cis*-regulation of RNA editing. We integrated high-throughput measurements of ADAR1 editing with computational analysis of more than 100 RNA sequence and structure features simultaneously. The CRISPR/Cas9 engineering allowed us to study the *cis*-regulation of RNA editing *in vivo* due to its minimal manipulation of the RNA editing process. Substrate-specific machine learning models that integrated diverse sequence and structural features explained over 71% of the variance in held-out variant test sets and enabled prediction of editing levels on unseen variants (**Fig. 5**). These models also provided insights into the contributions of different predictive features to ADAR editing (**Fig. 6**). Our approach is more biologically relevant compared to approaches using synthetic oligo library to mimic dsRNA substrates because RNA variants engineered by CRISPR/Cas9 are introduced to endogenous genomic loci. A lot of progress has been made in recent years in deciphering the splicing code^36–40^. However, the ADAR-mediated RNA editing code likely harbors more complex regulation via RNA secondary structure.

Our approach opens several new lines of inquiry. Measuring the structure of RNA variants in cells^41–43^ at native endogenous loci (vs. relying on predicted structure) would greatly enhance the information content. Further, while we focused on *cis* elements immediately adjacent to the editing site, longer-range interactions may be important for editing and can be investigated using our approach. Other factors that can be evaluated in the future include the regions in the RNA substrates that may be bound by the dsRNA binding domains (such as illustrated in **Fig. 5c**)^44^, long-range *cis* elements such as the editing inducer elements^45^, and *trans* regulators such as RNA binding proteins^46^. While our current results give rise to models with substantial predictive power for individual substrates, their generalizability remains low (**Supplementary Fig. 6**). This tendency to overfit will be mitigated by expanding the training set to include more RNA substrates, allowing the model to learn the shared properties of RNA substrates.

Several systems were recently developed to recruit ADAR enzymes to specific sites for site-directed RNA editing^23–29^, providing novel tools to study biological function and a safer and reversible alternative to gene therapy^23,47–49^. These RNA engineering approaches use antisense RNA oligoribonucleotides as guide RNAs (gRNAs) to recruit either engineered or endogenous ADAR enzymes. Currently, the RNA engineering methods utilizing ADAR enzymes usually use guide RNAs (gRNA) that perfectly duplex with the target region except for an A:C mismatch at the editing site^9–14^. Our results provide new possible designs to mimic the highly selective and efficient editing observed in some natural ADAR substrates with complex non-perfect duplex structures. Such features include relatively short 5’ (upstream) stem required for ADAR1^17^ compared to ADAR2 and specific non-stem 3’ (downstream) structure, where the internal loops at the 3’ likely contribute to the ADAR1 selectivity^22^. Notably, each of the three RNA substrates we tested has substrate-specific features that dictate the editing efficiency (**Fig. 5f**). This showcases that a screen of possible designs of gRNA would be a valuable and cost-effective strategy to learn the best features that lead to the most specific and efficient editing for each different target site. In this regard, our experimental methods and computational pipeline are readily applicable, identifying features important for editing and predicting editing levels around a particular site of interest. This will increase the repertoire of strategies that could lead to more precise and efficient editing of target adenosine sites in the transcriptome, which are highly desirable for safely correcting disease-related mutations at the RNA level.

Building on our work using the PREUSS pipeline, ADAR editing can be further investigated in larger scale and in different cell types, tissues and disease states to explore the full spectrum of *cis* regulation. Ultimately, establishing this “editing code” will help us better understand the underlying rules of RNA editing, which will facilitate efficient and precise transcriptome engineering for studying RNA biology and treating human disease.

## Methods

### Cell culture and transfection

HEK293T cells were cultured in Dulbecco’s modified Eagle medium (Life Technologies) supplemented with 10% FBS (Gibco, Thermo Fisher) and penicillin streptomycin (Life Technologies). Cells were maintained at 70%-90% confluency. One day before transfection, around 700,000 cells were split to 6 wells. The next day, 500 ng of Cas9-sgRNA construct in the p×330 backbone (https://www.addgene.org/42230/) was co-transfected with 500 ng of the DNA donor using lipofectamine 2000 (Invitrogen). Cells were maintained at 50%−90% confluency for 5 days.

### Design of the CRISPR/KI donor oligos

Two types of CRISPR knockin (KI) donors were designed in this study: the degenerate donor and the fixed donor. For the degenerate donor oligos, a single stranded DNA oligo was synthesized in which degenerate sequences were introduced at the interrogated regions. In the NEIL1 donor, −3 to −1 and +1 to +3 were interrogated and equal molar of 4 nucleotides were introduced at these positions during DNA synthesis. To avoid cutting by the Cas9, a point mutation was also introduced at the PAM sequence. In the TTYH2 donor, a 10 nt region in the editing complementary sequence was studied, and equal molar of C or T was introduced. The PAM sequence was also mutated along with a compensatory mutation to maintain the secondary structure.

For donors used for targeted mutagenesis, individual DNA sequences were designed to carry desired mutation(s). Briefly, a 15-20nt region around the target editing site and the corresponding region on the opposite strand were subject to mutagenesis. All possible nucleotide at any single position was tested, with exceptions where A-to-G mutation was avoided in the +1 and −1 nt of NEIL1 because it potentially becomes indistinguishable with A-to-I editing in RNA-seq results. Combination mutations at two positions were also designed, in which each of the positions is mutated to the nucleotide in the opposite strand to disrupt the original structure. In addition, individual donors with altered length for interrogation of specific features of the RNA substrate were included.

### Generation of the CRISPR/KI donor pool

For NEIL1 donors, 80mer oligos were purchased from IDT and pooled at equal molar ratio. Oligo pairs NEIL1_leftarm/NEIL1_rightarm, or asymmetrical labeled primer pairs NEIL1_leftarm_biotin/NEIL1_rightarm and NEIL1_leftarm/NEIL1_rightarm_biotin were used separately to add additional sequences to obtain 200mers in PCR reactions using Phusion polymerase. Around 400ul PCR products were purified using MinElute PCR purification kit (Qiagen) to obtain the dsDNA donor pool which was verified by agarose gel electrophoresis. For single-stranded donors, 100ul MyOne Streptavidin Dynabeads (Thermo Fisher) were added to the purified products that were amplified with asymmetrical biotin label and then the mixtures were denatured at 95 C for 10min and chilled on ice immediately. The unbound single stranded oligos were collected from the supernatant and then purified with column MinElute PCR Purification Kit (Qiagen) to obtain the ssDNA donors. For TTYH2 and AJUBA donors, 100 to 120-mer pooled oligos were purchased from Agilent and amplified using individual primers (Supplementary Table 6). Primers donor_F and donor_R of each target gene were used to specifically amplify the oligo library from the oligo chip. A second PCR using Donor_F_70 and Donor_R_70 was performed to elongate the homologous arms of each donor. The PCR products were purified using MinElute PCR purification kit (Qiagen) and used as dsDNA donors later. NEIL1_degenerate_donor and TTYH2_degenerate donors were synthesized as Ultramer by IDT and used directly in the transfection.

### Guide RNA design and cloning

Guide RNA was predicted by the web-based software CRISPR.mit.edu. The higher ranked guide RNA with a PAM sequence close to the interrogation region was selected. For the TTYH2 and AJUBA loci, different sets of guide RNAs were designed for the KI regions in two opposite strands. To construct guide RNA plasmids, two reverse complementary single stranded oligos with overhangs were synthesized by IDT and annealed on a thermocycler (Bio-Rad) before ligation to BbsI-linearized PX330 backbone. The ligation mix was transformed into Stbl3 chemical competent cells (Invitrogen) and single clones were sequence verified by Sanger sequencing.

### CRISPR mutagenesis and library construction

We used 600 ng single stranded oligo donor library or 1200 ng double stranded oligo donor library along with 500 ng guide RNA construct to co-transfect into 1 million HEK293T cells using lipofectamine 2000 (Invitrogen). For degenerate donor mediated knock in, 1ul of 10uM degenerate donor was used. 1uM L755507 (Sigma Aldrich) was added to the media one day after transfection to enhance the HDR efficiency as reported previously^50^. Two biological replicates were included for each assay. The transfected cells were grown for 5 days before they were seeded onto 10 cm dishes for an additional two days. 10% of the cells were harvested for genomic DNA using Quick-DNA kits (Zymo Research). The remaining cells were used for nuclear extraction using the Nuclear/Cytosolic Fractionation Kit (Cell Biolabs) following the manual. Nuclear RNA was purified from the nuclear extract using the Trizol method. Genomic DNA was removed from the RNA samples using the TURBO DNase (Thermo Fisher Scientific). RT was performed using SuperScript III kit (Thermo Fisher Scientific) and the gene specific primers. All RT products were used in total of 300 ul (50 uL × 6) PCR reaction with Phusion polymerase (Thermo Fisher Scientific) and gene specific primers with Fluidigm mmPCR adaptor sequences^51^. Genomic DNA library was amplified using similar approach except for the different primer set. All first round of PCR products were size selected on 1.5% Agarose gel and purified using Gel purification Kit (Qiagen). Diluted PCR product (1:50) was used in second round of PCR to add the Illumina sequencing adapter and individual barcode sequences using Fludigm_universal_F/fludigm_barcode_R^51^. The library was size selected and purified as in the previous step.

### Next-generation sequencing and data analysis

All libraries were sequenced on a NextSeq 550. NEIL1 libraries were sequenced for 75 cycles paired end and TTYH2 and AJUBA libraries 150 cycles paired end. To map the variants of the target gene, a reference genome was first built using the GMAP package, where designed mutations were included as SNPs. Briefly, GSNAP was used to detect variants with mismatches inside the interrogated region but not INDELs. The mapped reads were separated into individual variants based on the unique mutations carried in the region except for the editing site and RNA editing was called and measured for each variant as described in previous report. The indel variants were mapped individually. The z-score of each variant is calculated for each RNA library as:

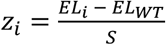; where 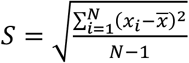; *EL* = *editing level*; *x*_*i*_ = *EL*_*i*_ − *EL*_*wT*_; 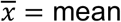 editing level for a given library.

### RNA secondary structure prediction

The sequence used for WT NEIL1, TTYH2 and AJUBA are shown in **Supplemental Table 7** and **Fig. 1b**. For AJUBA, we chose sequence for the region near the editing site by omitting 524 nt sequences in lieu of the full length (>800 bp). The secondary structures (with the minimum free energy) of the RNA variants for all three RNAs are calculated from the Vienna RNAfold 2.4.14 using default parameters^42^.

### Calculating the probability of forming wild type secondary structure

The probability of forming wild type-like RNA secondary structure was calculated with Vienna RNAfold version 2.1.9. The probability of forming the wild type secondary structure was calculated as:

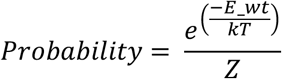

Where kT = 0.6 kcal at temperature T = 37 °C. Z is the unconstrained partition function (calculated with RNAfold −p). E_wt is the energy of the state with the wild type-like secondary structure, calculated using the following constraints in RNAfold, where ‘>’ indicates that the given base must be paired with a residue that comes before it (5’) in the sequence and ‘.’ indicates no constraint for the given base:

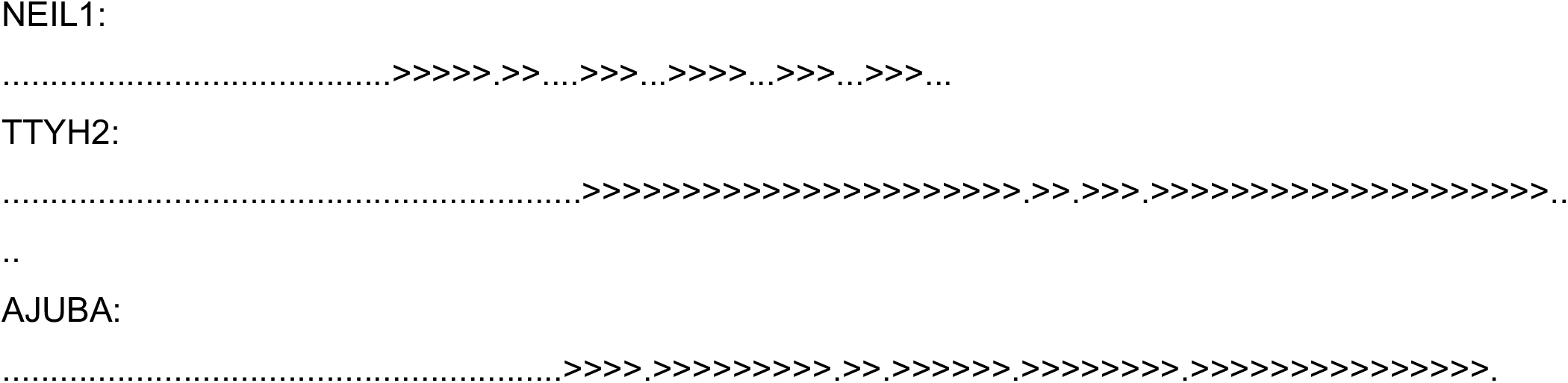

Z is the unconstrained partition function (calculated with RNAfold −p). An additional penalty was added if base pairs could not be formed in the core region of the RNA. T is 310.15 kelvin (37 °C) For NEIL1, the probability was divided by the number of non-canonical base pairs that would be formed in the core of the wild type secondary structure, according to the following secondary structure, to roughly account for the additional energetic penalty that these base pairs should incur. Such reference information used for each RNAs are shown below.

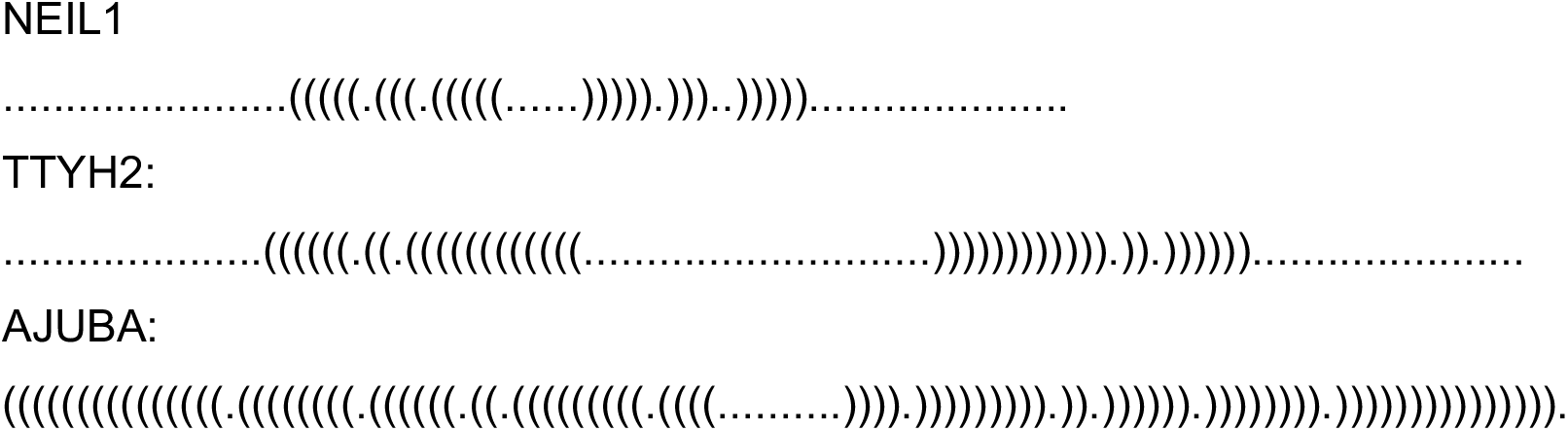

### Modeling the 3D structure of ADAR1 bound to NEIL1

A homology model of human ADAR1 deaminase was built using Phyre2^52^. The conformation of the core RNA residues (nucleotides corresponding to NEIL1 residues 30-39 and 44-53) was taken from the previously solved structure of human ADAR2 bound to double-stranded RNA (PDB ID: 5HP3). The RNA was positioned relative to the protein by aligning the previously solved ADAR2 structure (in complex with dsRNA) to the homology model of ADAR1, then copying the RNA coordinates from the ADAR2-dsRNA structure. Protein residues in the ADAR1 homology model that clashed with the RNA were removed (the final residues included in the model were: 823-973, 996-1003, and 1010-1223). The remaining NEIL1 RNA nucleotides were modeled using the Rosetta RNP-denovo method^53^ with the following command line:

rna_denovo -fasta fasta.txt -secstruct_file secstruct.txt -s ADAR1_homology_model_and_core_RNA_from_5hp3.pdb RNA_helix_1.pdb RNA_helix_2.pdb RNA_helix_3.pdb RNA_helix_4.pdb RNA_helix_5.pdb -new_fold_tree_initializer true-minimize_rna true -minimize_protein_sc false -out:file:silent build_full_wt_neil1.out -rna_protein_docking true - rnp_min_first false -rnp_pack_first false -cycles 10000 -rnp_high_res_cycles 0 -minimize_rounds 2 - nstruct 2000 -ignore_zero_occupancy false -convert_protein_CEN false -FA_low_res_rnp_scoring true-ramp_rnp_vdw true-dock_each_chunk_per_chain false-use_legacy_job_distributor true-no_filters

where RNA_helix_1.pdb, RNA_helix_2.pdb, etc. are ideal A-form helices for base-paired regions of Neil1. Possible placements of the double stranded RNA binding domains were visualized by aligning the previously solved structure of the ADAR2 dsRNA binding motif bound to dsRNA (PDB ID: 2L2K) to our model of NEIL1 bound to ADAR1.

### Machine learning models of RNA editing levels

All feature extraction and model training code is available to access on github: https://github.com/kundajelab/adar_editing

### Feature extraction

Boot-strapped and inferred structures for NEIL1, AJUBA, and TTYH2 were annotated with the bpRNA algorithm^30^. The bpRNA annotations were in turn utilized to extract structural and positional features for each variant. A feature matrix with structure-specific features, sequence-specific features, and stability-specific features (**Supplementary Table 1**) was engineered for each substrate and included a total of 122 features (**Supplementary Tables 2, 3, 4**).

### Model training

The XGBoost^44^ Python library (v. 0.81) was used to train gradient boosted regression trees to predict Adar editing levels from feature matrices described above. Training was performed both within-substrate and across substrates. The following approaches were utilized:

- Train on NEIL1, predict on NEIL1.
- Train on TTYH2, predict on TTYH2.
- Train on AJUBA, predict on AJUBA.
- Train on NEIL1 and TTYH2, predict on AJUBA.
- Train on NEIL1 and AJUBA, predict on TTYH2.
- Train on TTYH2 and AJUBA, predict on NEIL1.
- Train on TTYH2, AJUBA, NEIL1. Predict on TTYH2, AJUBA, NEIL1.
- Train on NEIL1, predict on AJUBA, TTYH2
- Train on TTYH2, prediction on NEIL1, AJUBA
- Train on AJUBA, predict on NEIL1, TTYH2

The dataset was randomly separated into 3 splits: training on 70% of variants, model validation on 15%, and testing on the remaining 15%. To avoid train/test contamination, base pair positions along the RNA molecules were assigned to one of the 3 splits (training, tuning, or test). All features associated with a given base pair position were assigned to the corresponding split. Any feature that was null or non-varying across all variants in a given training splits was removed from analysis. Any variant that had more than one mutation was included in the feature matrix twice -- the features in each entry for the variant were calculated specifically for one of the mutations.

XGBoost was trained for a maximum of 1000 iterations, with early stopping after 10 subsequent rounds with no reduction in root mean square error (RMSE) on the validation split. Default parameters were used.

In addition to models trained on computed feature values as indicated in **Supplementary Tables 2-4**, a separate set of models were also trained across substrates using ratios of numerical feature values relative to the wild type feature values. Ratios of feature values relative to the WT feature values within the same substrate were computed for the following feature set: ‘editing_value’, ‘free_energy’, ‘sim_nor_score’, ‘probability_active_conf’, ‘all_stem_length‘,‘site_length’, ‘site_length_internal_es’,‘site_length_internal_ecs’, ‘u_count’, ‘u_all_stem_length’, ‘u_hairpin_length’, ‘u1_distance’, ‘u1_length’, ‘u2_distance’,‘u2_length’, ‘u3_distance’,‘u3_length’, ‘d_count’,‘d_all_stem_length’, ‘d1_distance’,‘d1_length’, ‘d2_distance’,‘d2_length’, ‘d2_length_internal_ecs’, ‘d3_distance’, ‘d3_length’. Ratios were also calculated for substrate-specific features such as the length of hairpin vs stem vs bulge for the upstream and downstream structural features.

The R^2 value was calculated on the test set to determine the percent of total variance explained by the feature matrix^31^. Other metrics to measure model performance included:

- Spearman correlation from the scipy.stats Python library
- Pearson correlation the scipy.stats Python library
- Mean absolute error (MAE) from sklearn.metrics Python library
- Mean absolute percent error (MAPE)
- Root mean square error (RMSE) from sklearn.metrics Python library
- Area under the precision recall curve (auPRC) from sklearn.metrics Python library
- Area under the receiver operating characteristic (auROC) from sklearn.metrics Python library

### Feature importance analysis

Feature importance analysis was performed to identify the subset of features most informative in predicting Adar editing levels. The XGBoost “plot_importance” function was used to calculate the F score for each feature. The TreeSHAP algorithm^31^ was applied to interpret feature importance from the XGBoost model. SHAP summary values were computed for each feature as a measure of feature importance using the “shap_values” function within the “TreeExplainer” class. Pairwise interaction values from TreeShap were also calculated to identify highly correlated feature values.

SHAP values were applied to calculate the combined relative importance of feature subsets. Feature subsets (**Supplementary Table 1**) were defined as follows; some features were parts of multiple subsets:

- Structure features: stem length, free energy, probability of active conformation.
- Number of mutations in the variant.
- Mutation-specific sequence features: mutation position, mutation site reference allele, mutation site alternate allele, distance of mutation site from edited base.
- Mutation-specific structure features: bpRNA structure designation for the mutation site, bpRNA structure designation for the adjacent upstream site, bpRNA structure designation for the adjacent downstream site, boolean indication of whether or not the mutation is part of the same structure as the editing site.
- “Other” mutation-specific features: Type of mutation (indel, SNP), presence/absence of mutation (WT/ mutated) in the variant
- Editing site sequence features
- Editing site structure features
- Characterization of the 3 structural features upstream of the editing site
- Characterization of the 3 structural features downstream of the editing site

For each feature subset, the mean absolute SHAP values across variants were calculated. These were in turn summed across all features in the subset and compared to the total sum of mean absolute SHAP values across all features.

Overall feature rankings were computed by calculating the mean absolute value of SHAP values for each feature across the test set samples. These mean(|SHAP|) values were summed across all features and the percent contribution to the total was obtained for each feature. These percent contributions for each feature were averaged across substrates to determine features that were ranked as high importance consistently across all substrates.

## Data availability

The RNA-seq data are deposited in the following repository: Repository/DataBank Accession: GEO; AccessionID: GSE138860. Databank URL: http://www.ncbi.nlm.nih.gov/geo/. Bioinformatics codes for RNA editing call are available upon request. Codes for the PREUSS computational pipeline is available on GitHub URL: https://github.com/kundajelab/adar_editing.

## Acknowledgements

We thank members of Li, Kundaje, and Das laboratories, especially Dr. Patricia Deng, Dr. Anne L. Saprio, Dr. Shibin Hu, and Avanti Shrikumar for insightful discussions on the machine learning analysis, and Wipapat Kladwang and Dr. Joseph D. Yesselman for suggestions on RNA structure analysis. We also thank Lei Shi as a computational consultant. This work is supported by National Institutes of Health (NIH) (GM124215 and GM102484 to J.B.L), the Milton Safenowitz Postdoctoral Fellowship from the ALS Association (to T.S.) and the Stanford Bio-X Bowes Fellowship (to A.S.).

## Author contributions

X.L., T.S., G.R., and J.B.L conceived the work. X.L., A.S., T.S., A.K., J.B.L., I.J., and R.D. co-wrote the manuscript. T.S. designed and carried out the CRSIPR/Cas9 and RNA editing measurements. T.S. and Q.L. carried out the editing level analysis. X.L. and K.K. performed RNA and protein structure analysis. A.S., X.L., and A.K. developed the PREUSS computational pipeline.

## Legend for Supplementary Figures

**Supplementary Figure 1: Performance of targeted-mutagenesis RNA substrates. a)** Degenerate donor oligos are designed for a 10 nt region of the ECS in the TTYH2 substrate. The mutagenized region is highlighted in red and the editing site in blue. **b)** The distribution of editing level by the number of mutations from the results of the degenerate TTYH2-ECS library from (**a**). **c to f)** Reproducibility of the editing level measurement in different substrates. Each dot represents one designed variant. Editing level of each variant is compared. **c)** Two biological replicates for NEIL1. **d)** Two biological replicates targeting the TTYH2 edit strand. **e)** Two biological replicates targeting TTYH2 ECS. **f)** Two biological replicates targeting AJUBA edit stem. **g to h)** Donor type effect on CRISPR knock-in efficiency and RNA expression of NEIL1 variants. **i)** Abundance of designed NEIL1 variants detected from genomic DNA. Watson (sense) strand DNA oligo library or double stranded DNA library were used in the CRISPR mediated knock-in assay. Each dot represents one designed NEIL1 variant coverage in either experiment. **j to k)**. Editing level measurement reproducibility using different versions of NEIL1 donors, comparison between Watson strand donor and double stranded donor **(j)** and Watson strand donor and Crick strand donor **(k)**.

**Supplementary Figure 2: Editing levels from targeted-mutagenesis libraries. a)** Heatmap of editing levels from single- and double-mutations in the editing strand of TTYH2. **b)** Heatmap of editing levels from single mutations in the editing complementary sequence (ECS) of TTYH2. Editing level of WT TTYH2 is 0.31. **c)** Heatmap of editing levels from single- and double-mutations in the editing strand of AJUBA. Editing level of WT TTYH2 is 0.35. The z-score is calculated for each RNA library as described in Methods and the WT editing level z-score is 0. **d)** Comparing editing levels of variants with the mutation type. **e)** Comparing the normalized similarity score of the variants with mutation type. *, *P* < 0.05; ****, *P* < 0.0001; in t-test.

**Supplementary Figure 3: Selected examples of variants of NEIL1, TTYH2 and AJUBA. a-b)** Position-specific effects of TTYH2 **(a)** and AJUBA **(b)** single mutations. **c)** Effects of compound mutations that break the base pairing in the 5’ and 3’ structure of NEIL1 (WT NEIL1 editing level is 0.64). **d)** Comparing the editing levels of variants with different editing site structure. “Others” means the editing site A is located in a structure other than the 1:1 mismatch internal loop. The A:A mismatch was not presented in our RNA library.

**Supplementary Figure 4: Correlation of** *cis* **regulatory features to editing levels of all variants of NEIL1, TTYH2, and AJUBA. a)** Normalized Similarity Score. **b**) Free Energy. **c)** Probability of Active Conformation. **d)** All Stem Length. **e)** 5’ Stem Length.

**Supplementary Figure 5: Joint training and testing across substrates. a)**Test set XGBoost predictions for each substrate from a model trained jointly on NEIL1, TTYH2, and AJUBA. **b)** SHAP values for the 20 most important features driving test set predictions. Features are ranked in order of predictive importance from most important (top) to least important (bottom). **c)** SHAP values for the variants with highest and lowest editing levels for each substrate. **d)** Percent contribution to XGBoost test set predictions from each feature subset. Feature subset composition is defined in Supplementary Table 1. **e)** Spearman and Pearson correlation between observed and XGBoost-predicted editing levels for NEIL1, TTYH2, and AJUBA variants within the test set. **f)** Mean absolute error (MAE), mean absolute percent error (MAPE), and root mean square error (RMSE) for XGBoost predictions on test set variants of NEIL1, TTYH2, AJUBA. **g)** Area under the precision recall curve (auPRC) and area under the receiver operating characteristic curve (auROC) for test set variants of NEIL1, TTYH2, and AJUBA.

**Supplementary Figure 6: XGBoost model performance across training sets.** Spearman correlation between observed and predicted Adar editing levels in the held-out test set, as well as the percent of variance explained in the held-out test set, is illustrated for all combinations of training and test sets examined. Filled-in circles indicate that the particular substrate was included in a given training/test split.

## Legend for Supplementary Tables

**Supplementary Table 1: Feature engineering for machine learning.** Sheet 1: Features used with XGBoost for prediction of editing level; Sheet 2: List of of each feature sub-group with the features that comprise it.

**Supplementary Table 2: NEIL1 feature matrix**. Sheet 1 includes the degenerate variant feature values. Sheet 2 includes the computational variant feature values.

**Supplementary Table 3: TTYH2 feature matrix**. Sheet 1 includes the degenerate variant feature values. Sheet 2 includes the computational variant feature values.

**Supplementary Table 4: AJUBA feature matrix**. Computational variant feature values are shown.

**Supplementary Table 5: XGBoost prediction performance on the training, validation, and test splits.**

**Supplementary Table 6: Oligonucleotides for CRISPR/Cas9-mediated mutagenesis**. Sheet 1 is the guide RNA sequence. Sheet 2 and 3 contains sequences of donor oligos for CRISPR. Sheet 4 contains sequences of PCR primers.

**Supplementary Table 7: RNA sequence, mutation name, editing level and computationally predicted RNA secondary structure**. Sheet 1 is NEIL1. Sheet 2 is TTYH2. Sheet 3 AJUBA.

**Supplementary Table 8: Comparison of training jointly on all substrates with numerical feature values computed directly versus numerical feature value ratios relative to wild type**.

**Supplementary Table 9: Normalized SHAP and F1 scores for each feature used to train the substrate-specific AJUBA, NEIL1, and TTYH2 models**. Relative feature contributions are normalized to sum to 1 for the full set of features. A value of “NA” indicates that the feature was not used to train a given model due to a lack of training and/or validation examples for that feature/substrate combination.

## Notes

#### Summary of Updates

The sequence of the main figures was corrected in this revised version.

## References

1. Savva, Y.A., Rieder, L.E. & Reenan, R.A. The ADAR protein family. Genome Biol 13, 252 (2012).

2. Paz-Yaacov, N. et al. Adenosine-to-inosine RNA editing shapes transcriptome diversity in primates. Proc Natl Acad Sci U S A 107, 12174–9 (2010).

3. Nishikura, K. Functions and regulation of RNA editing by ADAR deaminases. Annual review of biochemistry 79, 321–49 (2010).

4. Li, J.B. & Church, G.M. Deciphering the functions and regulation of brain-enriched A-to-I RNA editing. Nat Neurosci 16, 1518–22 (2013).

5. Walkley, C.R. & Li, J.B. Rewriting the transcriptome: adenosine-to-inosine RNA editing by ADARs. Genome Biol 18, 205 (2017).

6. Melcher, T. et al. A mammalian RNA editing enzyme. Nature 379, 460–4 (1996).

7. Wang, Y., Zheng, Y. & Beal, P.A. Adenosine Deaminases That Act on RNA (ADARs). Enzymes 41, 215–268 (2017).

8. Hwang, T. et al. Dynamic regulation of RNA editing in human brain development and disease. Nat Neurosci 19, 1093–9 (2016).

9. Han, L. et al. The Genomic Landscape and Clinical Relevance of A-to-I RNA Editing in Human Cancers. Cancer Cell 28, 515–28 (2015).

10. Ramaswami, G. & Li, J.B. RADAR: a rigorously annotated database of A-to-I RNA editing. Nucleic Acids Res 42, D109–13 (2014).

11. Porath, H.T., Carmi, S. & Levanon, E.Y. A genome-wide map of hyper-edited RNA reveals numerous new sites. Nat Commun 5, 4726 (2014).

12. Bazak, L. et al. A-to-I RNA editing occurs at over a hundred million genomic sites, located in a majority of human genes. Genome Res 24, 365–76 (2014).

13. Tian, N. et al. A structural determinant required for RNA editing. Nucleic Acids Res 39, 5669–81 (2011).

14. Eggington, J.M., Greene, T. & Bass, B.L. Predicting sites of ADAR editing in double-stranded RNA. Nat Commun 2, 319 (2011).

15. Polson, A.G. & Bass, B.L. Preferential selection of adenosines for modification by double-stranded RNA adenosine deaminase. Embo J 13, 5701–11 (1994).

16. Lehmann, K.A. & Bass, B.L. Double-stranded RNA adenosine deaminases ADAR1 and ADAR2 have overlapping specificities. Biochemistry 39, 12875–84 (2000).

17. Wang, Y., Park, S. & Beal, P.A. Selective Recognition of RNA Substrates by ADAR Deaminase Domains. Biochemistry 57, 1640–1651 (2018).

18. Zhang, R., Deng, P., Jacobson, D. & Li, J.B. Evolutionary analysis reveals regulatory and functional landscape of coding and non-coding RNA editing. PLoS Genet 13, e1006563 (2017).

19. Sapiro, A.L., Deng, P., Zhang, R. & Li, J.B. Cis regulatory effects on A-to-I RNA editing in related Drosophila species. Cell Rep 11, 697–703 (2015).

20. Ramaswami, G. et al. Genetic mapping uncovers cis-regulatory landscape of RNA editing. Nat Commun 6, 8194 (2015).

21. Park, E. et al. Population and allelic variation of A-to-I RNA editing in human transcriptomes. Genome Biol 18, 143 (2017).

22. Lehmann, K.A. & Bass, B.L. The importance of internal loops within RNA substrates of ADAR1. J Mol Biol 291, 1–13 (1999).

23. Woolf, T.M., Chase, J.M. & Stinchcomb, D.T. Toward the therapeutic editing of mutated RNA sequences. Proc Natl Acad Sci U S A 92, 8298–302 (1995).

24. Stafforst, T. & Schneider, M.F. An RNA-deaminase conjugate selectively repairs point mutations. Angew Chem Int Ed Engl 51, 11166–9 (2012).

25. Montiel-Gonzalez, M.F., Vallecillo-Viejo, I., Yudowski, G.A. & Rosenthal, J.J. Correction of mutations within the cystic fibrosis transmembrane conductance regulator by site-directed RNA editing. Proc Natl Acad Sci U S A 110, 18285–90 (2013).

26. Wettengel, J., Reautschnig, P., Geisler, S., Kahle, P.J. & Stafforst, T. Harnessing human ADAR2 for RNA repair - Recoding a PINK1 mutation rescues mitophagy. Nucleic Acids Res 45, 2797–2808 (2017).

27. Vogel, P. et al. Efficient and precise editing of endogenous transcripts with SNAP-tagged ADARs. Nat Methods 15, 535–538 (2018).

28. Merkle, T. et al. Precise RNA editing by recruiting endogenous ADARs with antisense oligonucleotides. Nat Biotechnol 37, 133–138 (2019).

29. Qu, L. et al. Programmable RNA editing by recruiting endogenous ADAR using engineered RNAs. Nature Biotechnology (2019).

30. Findlay, G.M., Boyle, E.A., Hause, R.J., Klein, J.C. & Shendure, J. Saturation editing of genomic regions by multiplex homology-directed repair. Nature 513, 120–3 (2014).

31. Yeo, J., Goodman, R.A., Schirle, N.T., David, S.S. & Beal, P.A. RNA editing changes the lesion specificity for the DNA repair enzyme NEIL1. Proc Natl Acad Sci U S A 107, 20715–9 (2010).

32. Danaee, P. et al. bpRNA: large-scale automated annotation and analysis of RNA secondary structure. Nucleic Acids Res 46, 5381–5394 (2018).

33. Matthews, M.M. et al. Structures of human ADAR2 bound to dsRNA reveal base-flipping mechanism and basis for site selectivity. Nat Struct Mol Biol 23, 426–33 (2016).

34. Thomas, J.M. & Beal, P.A. How do ADARs bind RNA? New protein-RNA structures illuminate substrate recognition by the RNA editing ADARs. Bioessays 39(2017).

35. Lundberg, S.M. & Lee, S.-I. Consistent feature attribution for tree ensembles. in arXiv e-prints (2017).

36. Rosenberg, A.B., Patwardhan, R.P., Shendure, J. & Seelig, G. Learning the sequence determinants of alternative splicing from millions of random sequences. Cell 163, 698–711 (2015).

37. Xiong, H.Y. et al. RNA splicing. The human splicing code reveals new insights into the genetic determinants of disease. Science 347, 1254806 (2015).

38. Baeza-Centurion, P., Minana, B., Schmiedel, J.M., Valcarcel, J. & Lehner, B. Combinatorial Genetics Reveals a Scaling Law for the Effects of Mutations on Splicing. Cell 176, 549–563 e23 (2019).

39. Jaganathan, K. et al. Predicting Splicing from Primary Sequence with Deep Learning. Cell 176, 535–548 e24 (2019).

40. Cheung, R. et al. A Multiplexed Assay for Exon Recognition Reveals that an Unappreciated Fraction of Rare Genetic Variants Cause Large-Effect Splicing Disruptions. Mol Cell 73, 183–194 e8 (2019).

41. Rouskin, S., Zubradt, M., Washietl, S., Kellis, M. & Weissman, J.S. Genome-wide probing of RNA structure reveals active unfolding of mRNA structures in vivo. Nature 505, 701–5 (2014).

42. Ding, Y. et al. In vivo genome-wide profiling of RNA secondary structure reveals novel regulatory features. Nature 505, 696–700 (2014).

43. Zubradt, M. et al. DMS-MaPseq for genome-wide or targeted RNA structure probing in vivo. Nat Methods 14, 75–82 (2017).

44. Masliah, G., Barraud, P. & Allain, F.H. RNA recognition by double-stranded RNA binding domains: a matter of shape and sequence. Cell Mol Life Sci 70, 1875–95 (2013).

45. Daniel, C., Widmark, A., Rigardt, D. & Ohman, M. Editing inducer elements increases A-to-I editing efficiency in the mammalian transcriptome. Genome Biol 18, 195 (2017).

46. Freund, E.C. et al. Unbiased identification of *trans* trans regulators of ADAR and A-to-I RNA editing. bioRxiv, 631200 (2019).

47. Vogel, P. & Stafforst, T. Site-directed RNA editing with antagomir deaminases--a tool to study protein and RNA function. ChemMedChem 9, 2021–5 (2014).

48. Vogel, P. & Stafforst, T. Critical review on engineering deaminases for site-directed RNA editing. Curr Opin Biotechnol 55, 74–80 (2019).

49. Chen, G., Katrekar, D. & Mali, P. RNA-Guided Adenosine Deaminases: Advances and Challenges for Therapeutic RNA Editing. Biochemistry 58, 1947–1957 (2019).

50. Yu, C. et al. Small molecules enhance CRISPR genome editing in pluripotent stem cells. Cell Stem Cell 16, 142–7 (2015).

51. Zhang, R. et al. Quantifying RNA allelic ratios by microfluidic multiplex PCR and sequencing. Nat Methods 11, 51–4 (2014).

52. Kelley, L.A., Mezulis, S., Yates, C.M., Wass, M.N. & Sternberg, M.J. The Phyre2 web portal for protein modeling, prediction and analysis. Nat Protoc 10, 845–58 (2015).

53. Kappel, K. & Das, R. Sampling Native-like Structures of RNA-Protein Complexes through Rosetta Folding and Docking. Structure 27, 140–151.e5 (2019).

