## Supplementary figures and images for "Learning *cis*-regulatory principles of ADAR-based RNA editing from CRISPR-mediated mutagenesis"

### Supplemental Figures 1-6

#### Supp_Figure 1

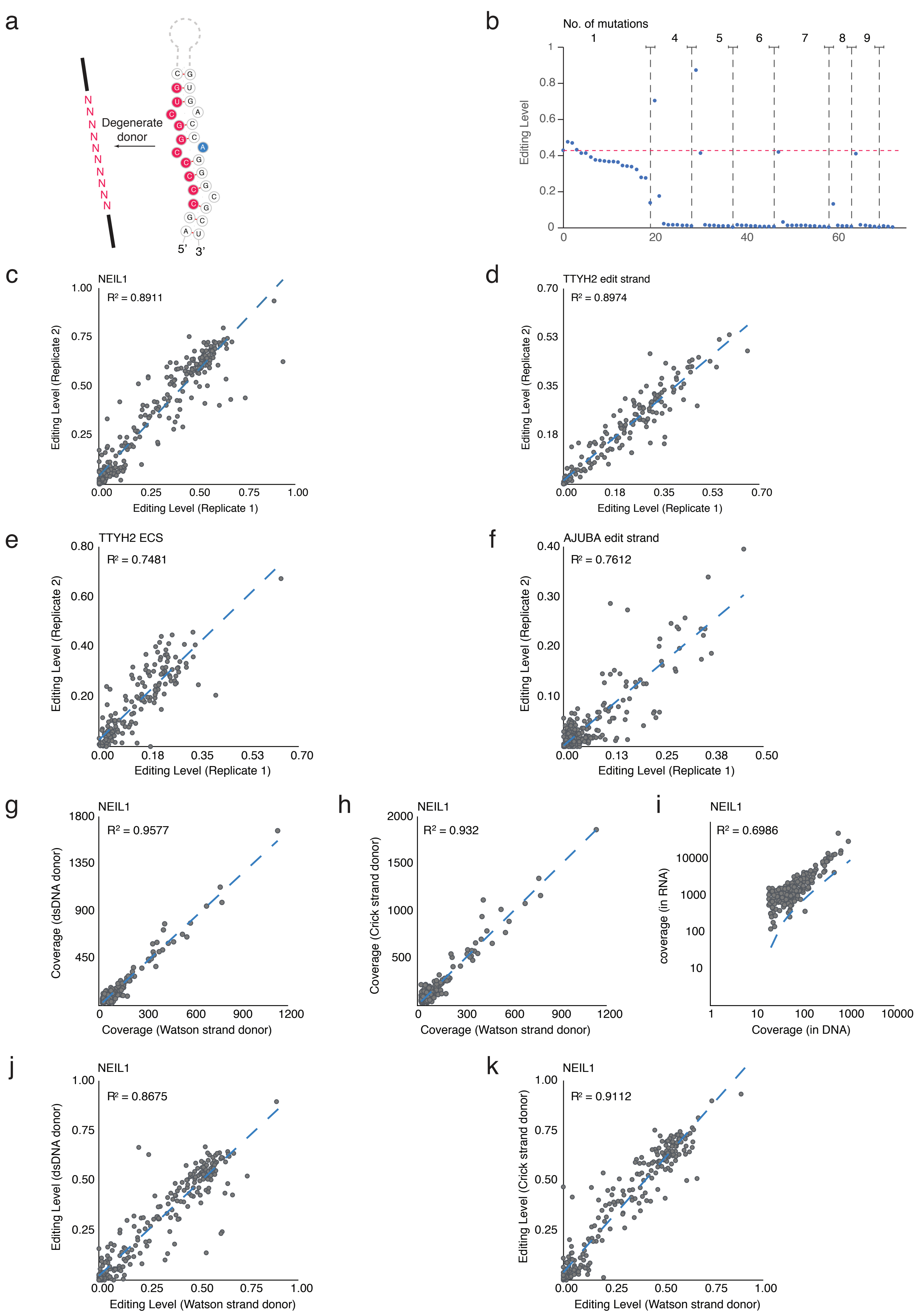

#### Supp_Figure 2

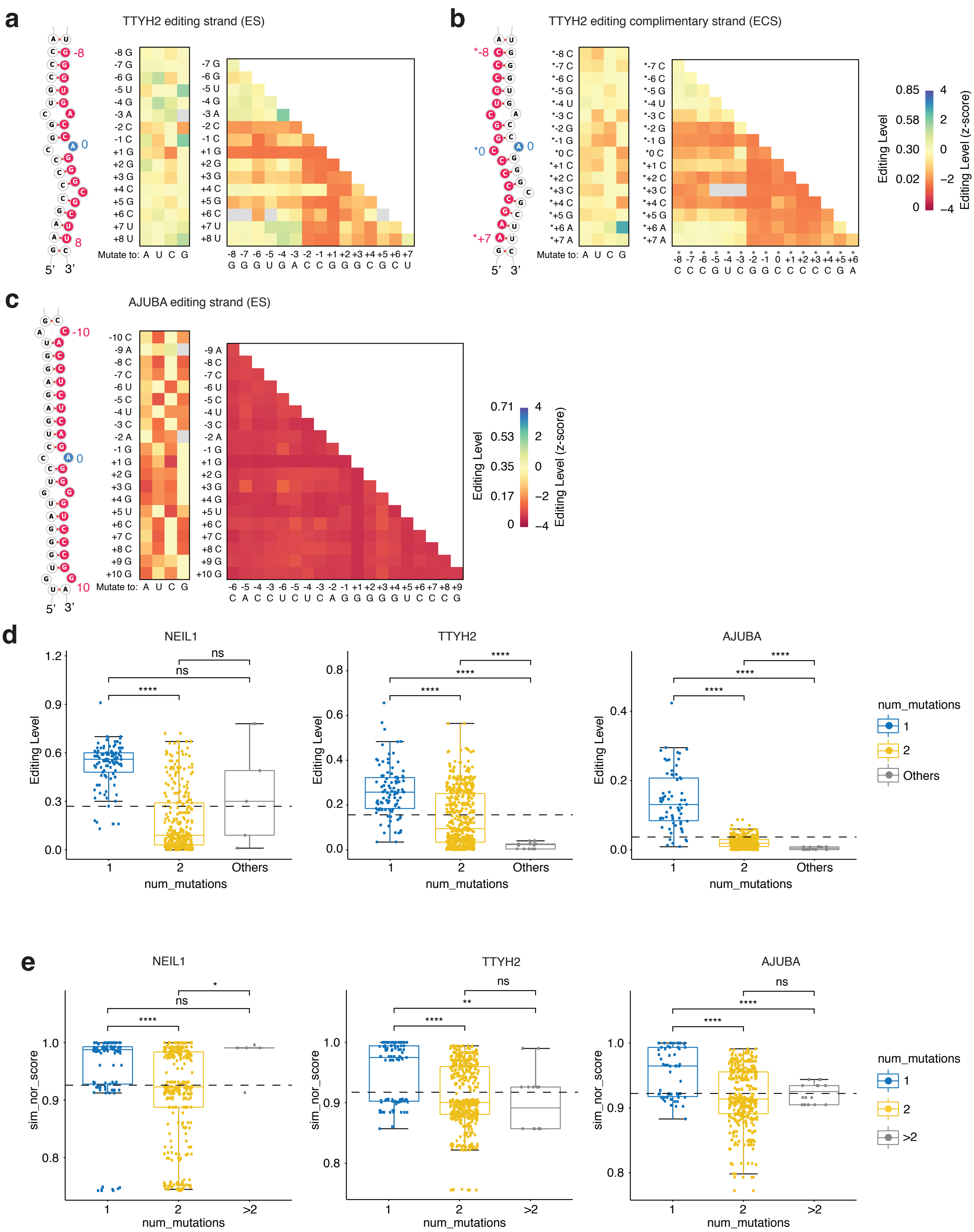

#### Supp_Figure 3

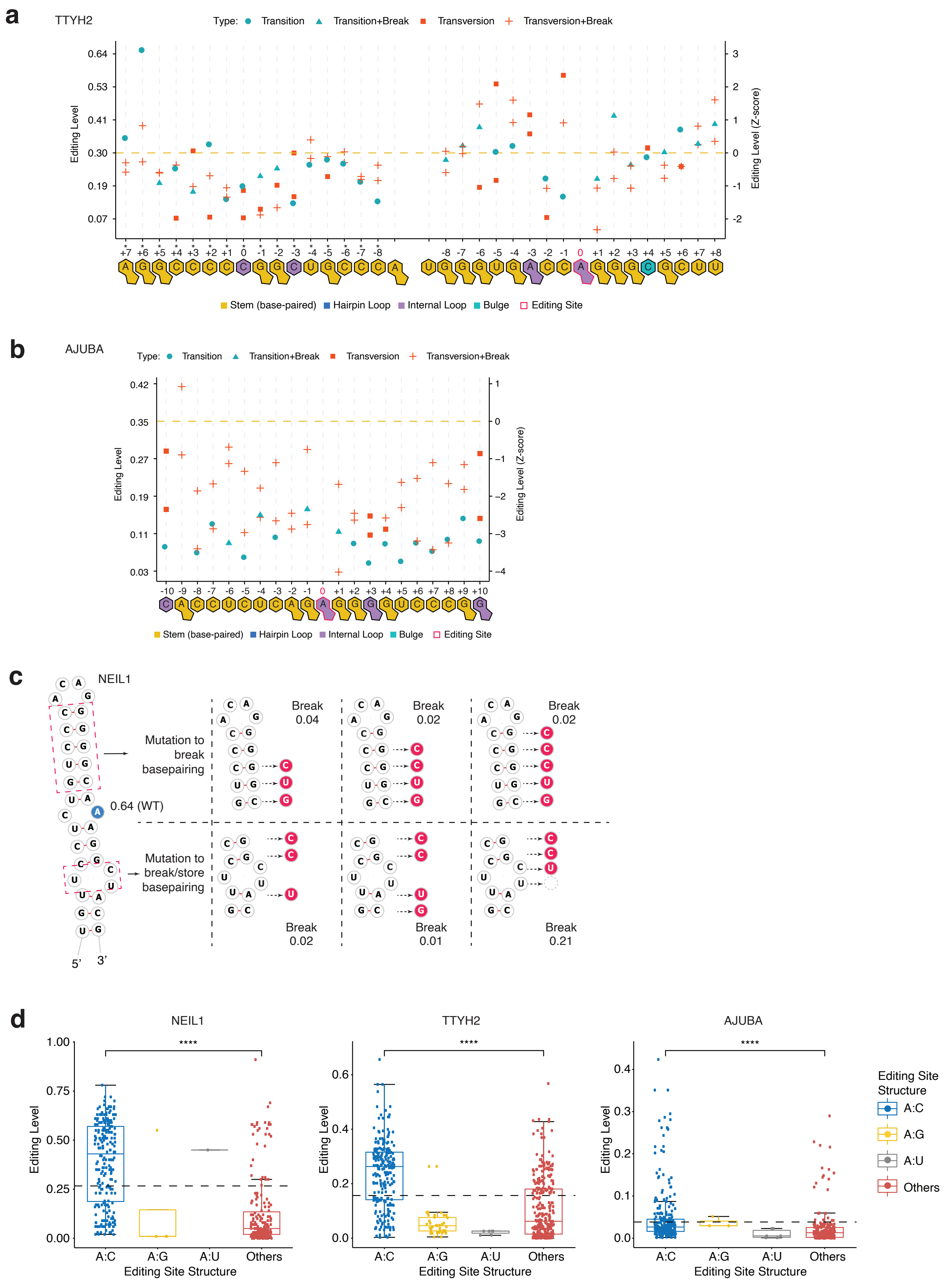

#### Supp_Figure 4

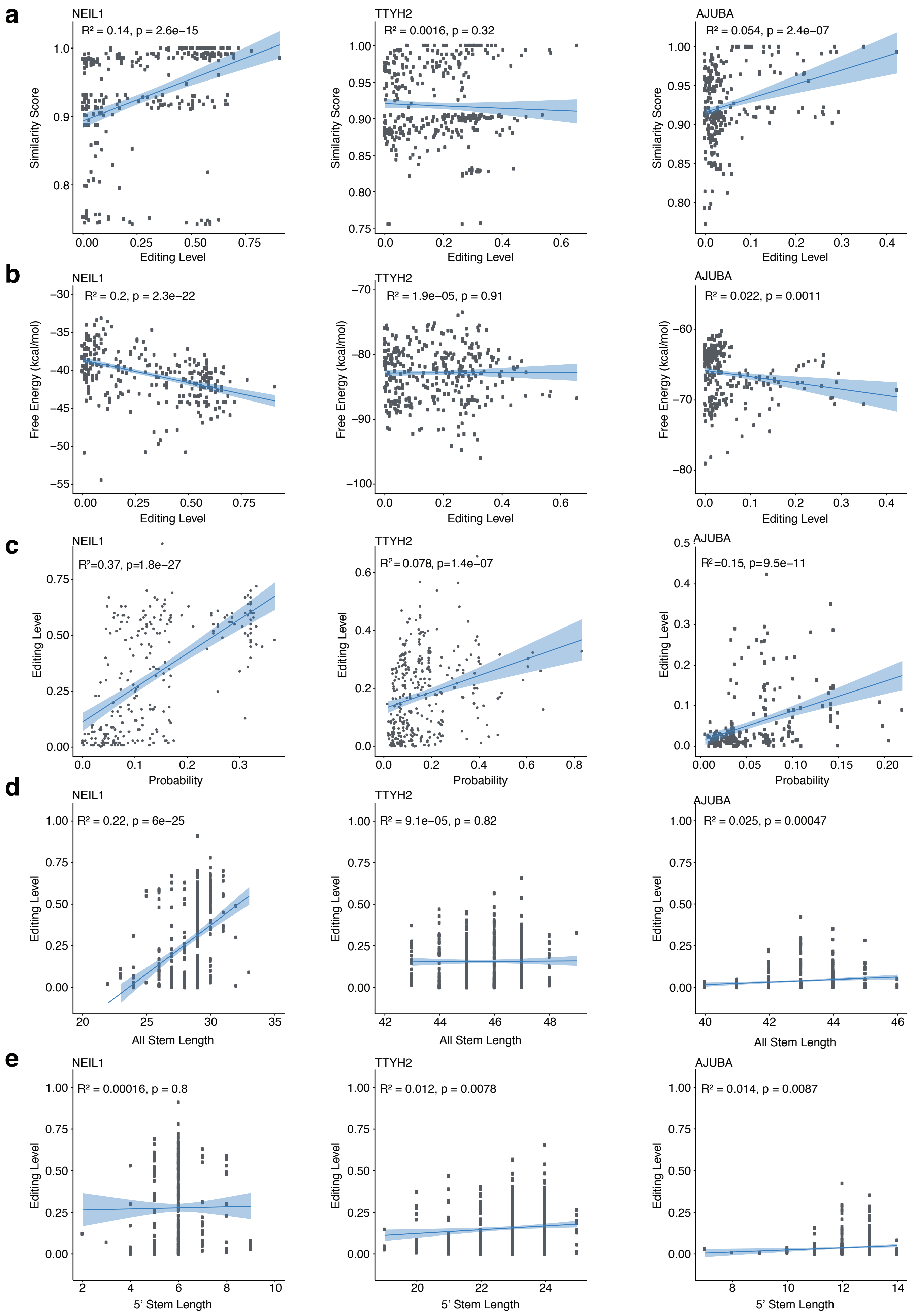

#### Supp_Figure 5

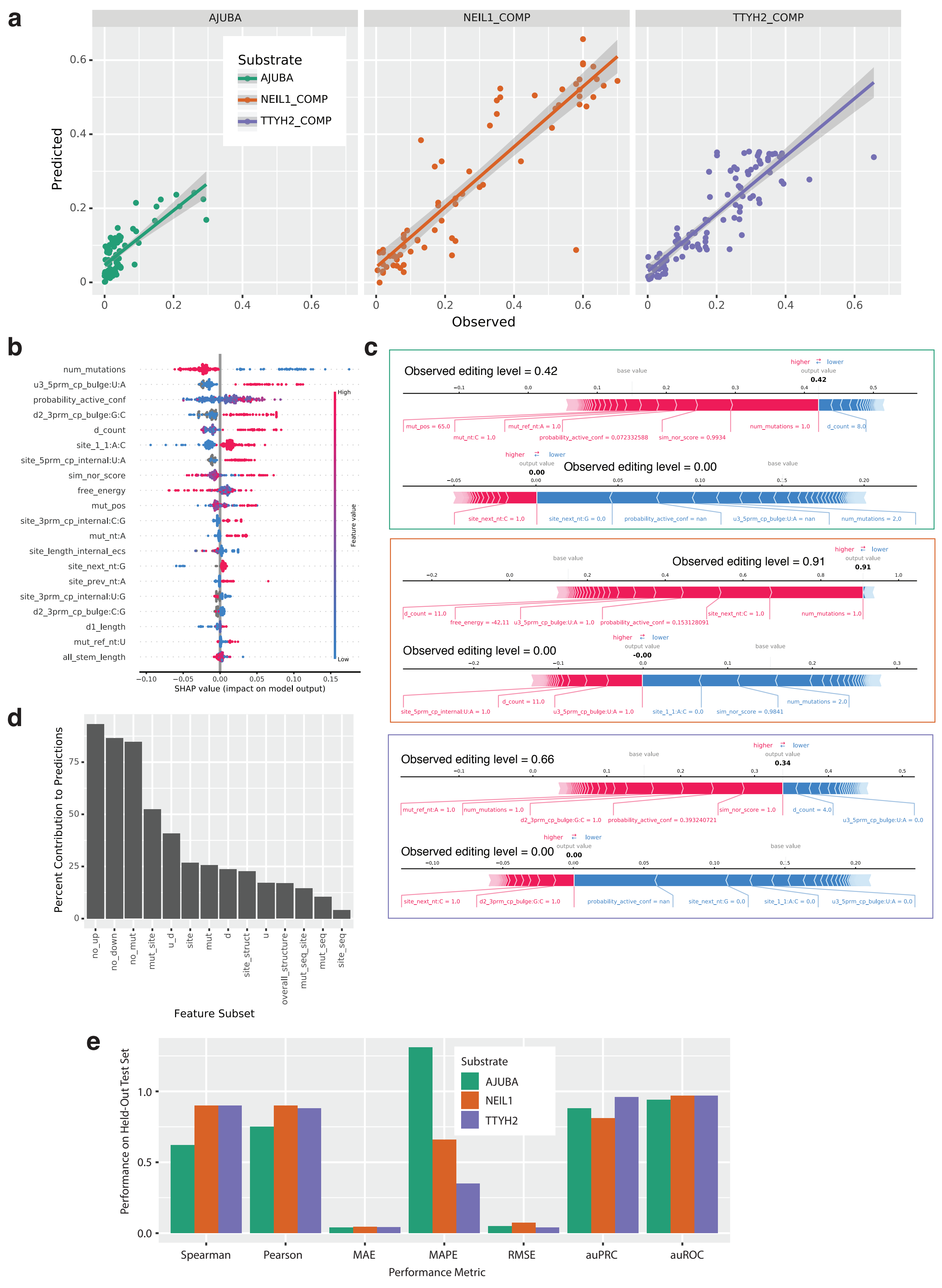

#### Supp_Figure 6

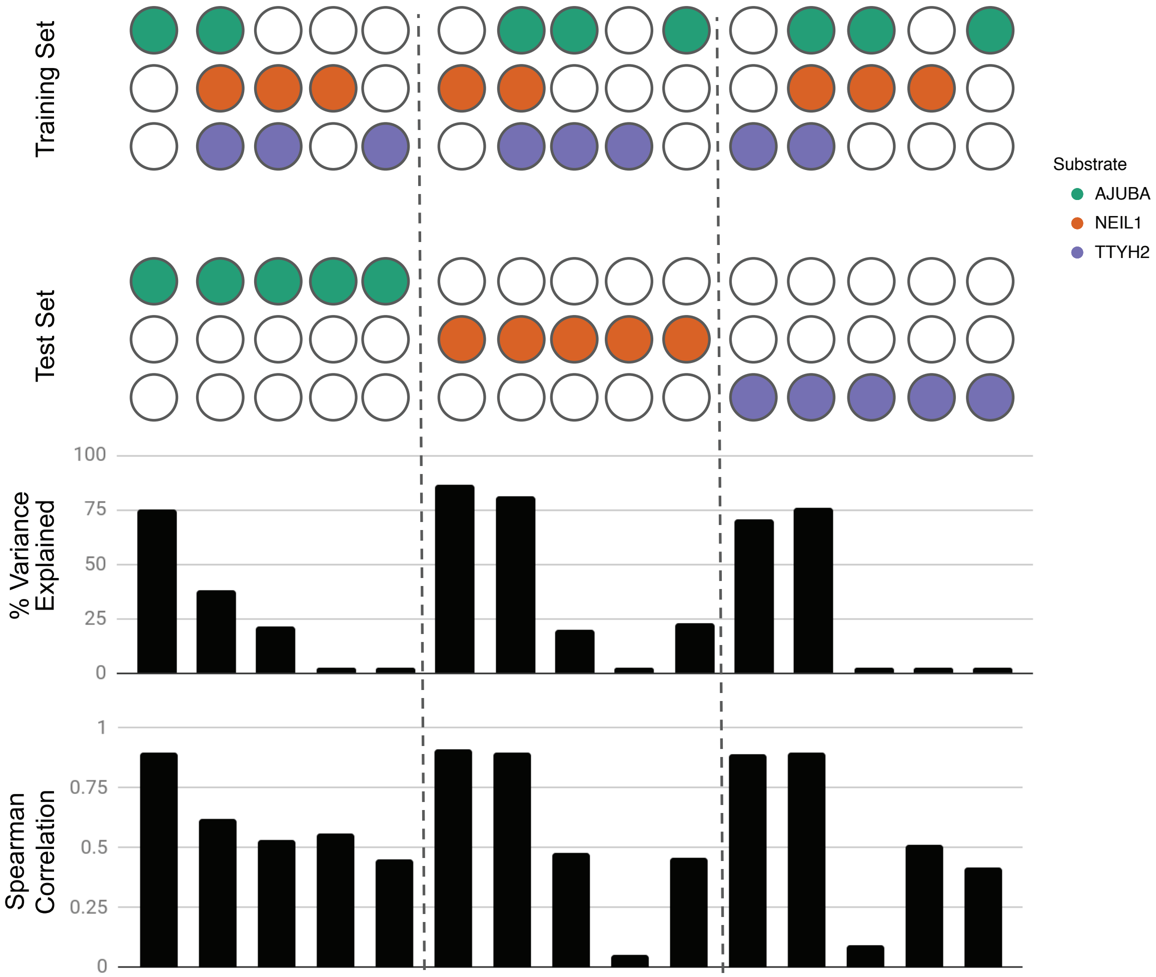
